# Breast tumors escape endocrine therapy by ER-independent mechanisms triggered by the coordinated activities of HER2/HER3 and deacetylated FOXA1

**DOI:** 10.1101/2020.09.22.308569

**Authors:** Siv Gilfillan, Shixiong Wang, Madhumohan R. Katika, Jens Henrik Norum, Helga Bergholtz, Elisa Fiorito, Siri Nordhagen, Yogita Sharma, Sachin Singh, Venkata S. Somisety, Anne-Marthe Fosdahl, Silje Nord, Olav Engebraaten, Ole Christian Lingjaerde, Anne-Lise Børresen-Dale, Kristine Kleivi Sahlberg, Therese Sørlie, Meritxell Bellet, Sandra Lopez-Aviles, Antoni Hurtado

## Abstract

Hormone-resistance in ER positive breast cancer is associated with high HER2 activity. Yet, the interplay between HER2 and FOXA1 in hormone resistant tumors is not elucidated. Now, we demonstrate that hormone resistant tumors have increased HER2 expression and that FOXA1 mediates the signals of HER2/3 in an Estrogen Receptor independent manner. Our *in vitro* and *in vivo* experiments reveal that HER2/HER3 triggers FOXA1 binding at chromatin regions of ER-regulated genes associated with poor prognosis, facilitating their expression and leading to ER-independent growth. Furthermore, our study supports that FOXA1 acetylated by the acetyltransferase EP300 is retained at ER chromatin regions, which enables ER function. By contrast, HER2/3 activation hinders FOXA1 acetylation and facilitates FOXA1 binding at non-ER interacting regions enriched towards poor prognosis genes. Moreover, FOXA1 deacetylation confers insensitivity to anti-ER drugs inhibitory effect in ER positive cells. These results elucidate how post-translational modifications of FOXA1 control transcription independently of ER in hormone-resistant tumors with enhanced HER2/3 signaling.

## INTRODUCTION

Luminal subtypes are the most frequent breast cancers and they express Estrogen Receptor alpha (ER) (1). Estrogen signaling exerts its growth-promoting effects by inducing ER binding to many chromatin sites (2), which leads to altered expression of coding and non-coding RNAs (3). Treatment with hormone therapies inhibits the function of ER and reduces tumor growth and improves survival. ER expression serves as an important diagnostic and predictive marker in breast cancer (4). Patients with ER positive and HER2 (Human Epidermal Growth Factor Receptor Factor 2) negative tumors show the best responses to endocrine agents. However, resistance to these agents has become a major clinical obstacle. Clinical studies have shown that recurrence on adjuvant endocrine therapy occurs in approximately 10–15% of patients with early stage ER-positive breast cancer within 5 years (5) and recurrence rates are as high as 30% by 15 years (6). Potential mechanisms of resistance to endocrine therapies have been identified, often involving enhanced growth factor signaling, mutations in ER and changes in the expression or action of ER. One common mechanism of resistance involves the loss of estrogen-ER regulation by members of the Human Epidermal Growth Factor Receptors (HER) family (7). HER members can activate the downstream MAPK/ERK and PI3K/AKT/mTOR pathways (8,9), which results in phosphorylation of ER (10). ER phosphorylation is associated with estrogen-independent ER transcriptional activity (11) and poor prognosis (12). By contrast, it is suggested that hormone resistant cancers may possess redundant survival and proliferation pathways, one mediated by ER and another mediated by HER family members (13). Despite years of extensive studies on the interplay between HER signaling and ER, the precise mechanism of their interaction is still unknown.

FOXA1, a transcription factor belonging to the Forkhead family, is a pioneer factor required for the expression of a majority of ER-regulated genes in the initiation and/or progression of luminal breast cancers (14). Furthermore, FOXA1 is still expressed in hormone-therapy resistant tumors (15). Previously it was suggested that FOXA1 might mediate the actions of HER2 signaling (16), but the clinical and biological significance of that interplay remains unknown. Moreover, and very recently, it has been described that PI3Kα inhibition mediates an open chromatin state at the ER target loci in breast cancer models (17). In our study we have identified that metastases from patients that relapsed on hormone-therapy showed increased levels of HER2 compared to the primary tumor from the same patient. Furthermore, we have also investigated the crosstalk between HER2 signaling and the transcription factor FOXA1. Our results pinpoint a new role for FOXA1 in breast cancers as a mediator of HER2/HER3 signaling and demonstrate that enhanced activity of this signaling pathway in luminal tumors confers ER-independent growth. Finally, we also investigated the mechanism by which FOXA1 controls growth of luminal tumors when ER is inhibited and HER2/3 signaling is triggered.

## MATHERIAL AND METHODS

### Cell culture

Cell lines were obtained from American Type Culture Collection (ATCC, Manassas, VA). BT474, MCF7-HER2, MCF-7 and MDA-MB-453high cell lines were cultured in DMEM (4.5 g/l glucose).

### Plasmids

HA-tagged FOXA1 was subcloned into pCI-neo (E1841, Promega, Madison, WI, USA) expression vector for transient transfection. Two lysines in Wing 1 (K237 and 240) and 3 lysines in Wing 2 (264, 267, 270) were mutated into arginines respectively (WD1R and WD2R) or all mutated (WD12R). The mutagenesis was carried out with QuikChange Lightning Site-Directed Mutagenesis Kit (Agilent, Santa Clara, CA,USA).

### Chromatin Immunoprecipitation (ChIP) and drug treatment

FOXA1 genomic regions were identified by using the cross-linking (X)-ChIP protocol as described previously (18). Cells were treated with anti HER2 drugs (Trastuzumab, 20⍰/ml; Lapatinib, 1⍰M), for 48 h and fixed with 1% formaldehyde. Chromatin was incubated with Chip grade FOXA1 antibodies (5⍰g of antibodies Abcam ab5089 and ab23738) and Protein A&G Agarose Beads (Life technologies). Library preparation for sequencing was done following the instructions of TruSeq DNA sample preparation kit from Illumina or MicroPlex Library preparation kit from Diagenode.

### Formaldehyde-Assisted Isolation of Regulatory Elements (FAIRE)

MCF-7 cells were grown in hormone-depleted media and low serum (2%) and treated with vehicle (DMSO) or Heregulin (25 ng/ml) for 1h. FAIRE experiment was performed as previously described (19).

### Transfection

Cells were seeded in 6 well plate to be 50% confluent upon transfection. Cells were transfected with siRNA targeting FOXA1 (ON-TARGET J-010319-05-0005, Thermo Fisher Scientific) and siControl Non-targeting (siNT) (SI03650318 from Promega) using Lipfectamine RNAiMax (Life technologies) to a final concentration of 55 nM. MCF-7 FOXA1 inducible expression stable cells were reverse tranfected with siRNA targeting EP300 (sc-29431, Santa Cruz) and siControl with the same method to siFOXA1 at a final concentration of 10nM.

For the transient transfection of FOXA1-WT and acetylation mutants, MCF-7 cells were transfected with Lipofectine 3000 (Invitrogen) following the manufacturer’s protocol.

### ChIP and FAIRE sequencing data Analyses

Reads generated by the genome analyzer were aligned against the human genome using Bowtie 2 software (http://bowtie-bio.sourceforge.net/bowtie2/index.shtml) with default parameters.

### Motif analyses

We used HOMER software for motif discovery identify ERE, FOXA1, AP2Gamma and PBX1 in the [–200 bp, +200 bp] window around the peaks identified in the FOXA1 ChIP sequencing data from MCF-7 and BT474 cells (20). Enrichment p-values reported by HOMER are assumed significant when p < 1e-50. The motif matrices were retrieved from the JASPAR database (21).

### RNA isolation and quality control

After siFOXA1 transfection, cells were isolated and the culture medium was removed after the centrifugation of the cell suspension. Total RNA was isolated with the total RNA isolation kit according to the manufacturer’s protocol (Life Technologies). NanoDrop 2000 assessed RNA yield.

### Microarray hybridization and data normalization

Total RNA (300 ng) was used with the Illumina TotalPrep Amplification Kit (Ambion) and hybridized to HumanHT-12 v4 Expression BeadChips (Illumina) enabling profiling of >47,000 transcripts.

### Microarray data analysis

Raw intensity values were exported from GenomeStudio® software (version 1.1.1) for data processing. Data quality and sample relations were assessed using the Bioconductor lumi package. Probes with a detection p-value less than 0.05 were considered present.

### Hierarchical clustering

Unsupervised hierarchical clustering was performed with the publicly available programs Cluster (uncentered correlation; average linkage clustering) and Tree view (22).

### Selection of differentially expressed genes

Genes were selected with Student’s *T*-test (FDR-value < 0.05 and P values <0.01). For this we used 2 log values of treatment vs. controls.

### Pathway analysis

Functional interpretation of differentially expressed genes for each drug treatment was done using Ingenuity software (Ingenuity, Qiagen).

### Gene set enrichment analysis (GSEA)

GSEA software (version 2.0.14) was used to detect the differential expression of biologically relevant gene sets. Gene sets with a p value ≤0.05 and a false discovery rate (FDR) ≤0.25 were considered significantly affected (23). GSEA sources: Biocarta-2 (http://www.biocarta.com/), KEGG, Reactome, Oncogenic signatures (MSigDB database v5.0) and poor prognosis gene signatures associated with poor prognosis in ER positive tumors or metastatic signature in breast cancer (15,24).

### Analysis of mRNA expression in breast cancer sections

Total RNA (150 ng) was shipped to the NanoString nCounter® Human mRNA Expression Assay analysis. RNA was incubated in the presence of mRNA specific probes. To account for minor differences in hybridization and purification efficiencies raw data was adjusted using a technical normalization factor calculated from six internal positive spike controls present in each reaction. Background hybridization was corrected by deducting the negative control mean plus two standard deviations calculated from eight negative controls.

### Cell proliferation assay

Cells were seeded at equal confluence and the effect of growth factor, drugs, siRNA or overexpression of FOXA1 on cell proliferation were measured by live cell imaging Incucyte (Essen BioScience). BT474, TAMR and MCF-7 cells were transfected with siFOXA1 or siNT oligonucleotides as described above and 48 h of post transfection treated with growth factors (EGF 100ng/ml and Heregulin 25ng/ml).

### Cell migration assay

MDA-MB-453 FOXA1 overexpressing cells were plated in ImageLock 96-well cell migration plate (Essen Bioscience). After confluence cells were treated with vehicle or 30ng/ml doxycycline to allow the overexpression of FOXA1. Then, scratch was made with a 96-pin WouldMaker (Essen Bioscience), and cells were treated with vehicle, EGF (100ng/ml) or heregulin (25ng/ml), with or without doxycycline. Relative wound density was monitored for total of 24 hours.

### Western blots

Protein lysate was resolved using precast SDS-PAGE gels, and transferred to PVDF membrane. Blots were blocked and incubated overnight at 4°C with primary antibodies. Antibodies used were: ERα (sc-543), p300 (sc-585), and ERBB3 (sc-285) from Santa Cruz Biotechnologies, HA (16B12) from Covance, FOXA1 (ab55178), ErbB2/Her2 (ab16901) and histone H3 (ab1791) from Abcam. Following antibodies from Cell Signaling Technology: RPL13A (2765), β-Actin (4970S), Phospho-HER2/ErbB2 (Tyr1221/1222) (2243), Acetylated-Lysine (Ac-K2-100) (9814).

### Chromatin fractionation

MCF-7, BT474 cells were grown in the same manner as described in the previous sections. Cells were scraped in cold PBS containing proteinase and phosphatase inhibitors (Thermo Scientific). Each cell pellet was incubated for 10 min in 200 μl of cold buffer A (10 mM Hepes [pH 7.9], 10 mM KCl, 1.5 mM MgCl_2_, 0.34 M sucrose, 10 % glycerol, 1 mM DTT, protease and phosphatase inhibitors (Thermo Scientific) supplemented with 0,1 % Triton X-100. The samples were subjected to low-speed centrifugation (4 min, 1,300g, 4 °C) to collect the nuclei. Pelleted nuclei were lysed in 200 μl of buffer B (3 mM EDTA, 0.2 mM EGTA, 1 mM DTT, protease and phosphatase inhibitors (Thermo Scientific), and kept on ice 30 min. Then, pellets were resuspended in 200 μl of buffer B and sonicated for 30 s (Bioruptor, Diagenode) to shear DNA. Chromatin was pelleted by centrifugation (5 min, 16,000g, 4 °C) and resuspended in 4× loading dye (Life Technologies GmbH).

### Patient derived xenografts (PDX) models of breast cancer and volume calculation

The serially transplantable luminal-like PDX model was established by implanting tumor tissue (2-3 mm) in SCID mice as previously described (25). Mouse with tumors maximally 1cm^3^ of volume were sacrificed and 1-2mm^3^ pieces of tumor tissues were directly transplanted in the mammary fat pad number 4 on both sides of 4-8 weeks old female NOD/SCID interleukin-2 receptor gamma chain null (Il2rg^-/-^) (NSG) mice. In experimental mice, when the tumor volume was 100-500 mm^3^ the administration of heregulin (5μg/day), EGF (10μg/day) or NaCl was initiated by implanting micro-osmotic pumps (Alzet) subcutaneously under general anesthesia. The osmotic pumps were set for drug delivery for up to 28 days. To the indicated animals, fulvestrant (Faslodex^®^, 5mg/mouse), trastuzumab (Herceptin^®^ 0,05mg/mouse) and/or NaCl was administered by subcutaneous injections twice weekly either as single agents or in combinations.

Tumor diameters (d_min_ and d_max_) were measured using a caliper and the tumor volume calculated using the formula d_min_^2^ x d_max_ x π/6. All further statistic analyses were conducted with tumor volumes relative to the corresponding tumor volume at time of pump insertion and treatment initiation. Linear regression and smoothing spline models were fitted to the relative tumor volumes for each of the ten treatment groups (n=5 mice per group). The spline model curves confirmed that the linear regression model satisfactory explained the tumor growth during the treatment period. All relative tumor volumes, regression model curves and splines. A two-sided Student’s t-test was performed to statistically test whether the slopes of the regression curves were significantly different from 0. Based on the linear regression model, the tumor volume at day 20 after pump insertion was calculated for each group.

## RESULTS

### HER2 expression is increased in metastases from hormone-resistant patients and controls FOXA1 binding to the chromatin

It is accepted that HER2 overexpression confers intrinsic resistance to hormonal treatment. Hence, in this study we first aimed to analyze the expression of FOXA1, HER2 and ER in patients with low-grade HER2 negative tumors who initially responded to anti-ER therapy but eventually relapsed (Supplementary table 1). We also analyzed metastases from the same patients with relapse. Moreover, primary tumors from patients who responded to treatment were also investigated. The mRNA and protein levels of HER2 were significantly increased in metastases compared to primary tumor biopsies of the same patient (Figure 1A and B, Supplementary Figure S1). Furthermore, ER or FOXA1 mRNA levels were not substantially affected (Figure 1A). These results suggested that an increase of HER2 might be the cause of resistance in these patients. Previously, it was suggested that FOXA1 might be regulated by HER2 signaling (26). Hence, we aimed to investigate whether FOXA1 might be mediating the actions of HER2 signaling. First, we analyzed FOXA1 chromatin interactions using ChIP sequencing (ChIP-seq) in HER2-positive (herein, HER2-high) and HER2-negative (herein, HER2-low) breast cancer cell lines. FOXA1 binding interactions were called using MACS (27), and HER2-low (MCF-7) and HER2-high (BT474) cell lines were examined. We found 75,765 FOXA1 binding events in MCF-7 and 124,930 in BT474 cells (Figure 1C). The comparison of FOXA1 binding between the two cell lines suggests that HER2 signaling pathway could influence the differential FOXA1-binding events observed in BT474 cells (56% unique sites). Interestingly, the genomic distribution analysis of FOXA1 sites revealed an enrichment of FOXA1 binding sites towards gene body and promoter at BT474 sites (unique and shared with MCF-7) compared to unique sites at MCF-7. The increase of FOXA1 binding at promoter was even higher in BT474 unique sites (Supplementary Figure S2A). We hypothesized that increased HER2 signaling might reprogram FOXA1 binding. To test the hypothesis we used MCF-7 cells overexpressing HER2 and performed FOXA1 ChIP-seq. We identified 122,301 FOXA1 binding events in MCF7-HER2 cells and the comparison with BT474 showed high overlap (62% for MCF7-HER2; 60% for BT474) (Supplementary Figure S2A). Intriguingly, the signal intensity of FOXA1 binding in MCF7-HER2 cells at chromatin regions exclusive for BT474 was significantly superior to the signal detected in MCF-7 cells (Figure 1D). We also compared MCF-7 cells with different levels of HER2 and confirmed that the increased expression of HER2 correlated with an increase in FOXA1 binding (54% unique sites in MCF7-HER2) (Supplementary Figure S2B). Next, we tested FOXA1-chromatin interactions on a genome-wide scale in the breast cancer cell lines treated with Trastuzumab, a monoclonal antibody that targets HER2. We also analyzed the signal intensity of FOXA1 chromatin interaction in control and treated cells (Figure 1E). The signal intensity of FOXA1 sites was significantly dampened by HER2 inhibition in all the cell lines tested, despite the fact that the amount of FOXA1 protein was not influenced (Supplementary Figure S2C). Surprisingly, FOXA1 binding was also reduced in HER2-low breast cancer cells (Figure 1E) when HER2 was inhibited (Supplementary Figure S2D). These data confirmed that HER2 positively regulates the binding of FOXA1 even in cells displaying low levels of HER2. Moreover these results support that FOXA1 might be reprogrammed in breast tumors with increased HER2 activity.

**Figure 1.**
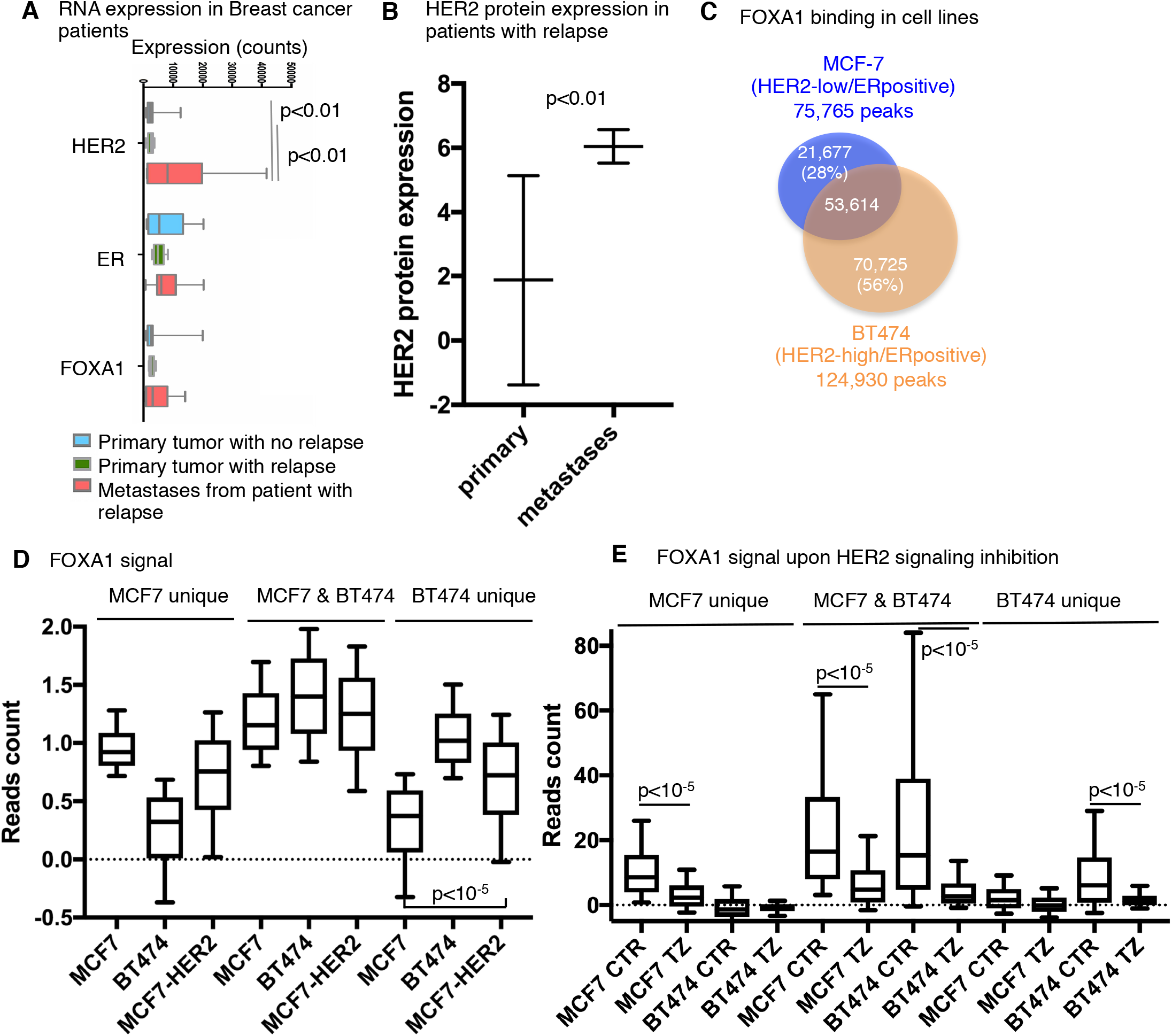
Differential binding of FOXA1 in breast cancer cells is perturbed by HER2 signaling inhibitors. (**A**) Box-plot indicating the expression of HER2, ER and FOXA1 in groups of ER positive patients from primary tumors from patients without relapse (n=10) and paired samples of metastases (n=5) and primary tumors from patients with relapse (n=5). Expression of mRNA was measured by using Nanostring techonology from formalin embedded samples. Wilcoxon rank-sum test was used to test any statistical difference between samples. (**B**) Box-plot indicating the protein expression of HER2 (expressed in logarithmic scale) in paired samples of metastases (n=3) and primary tumors from patients with relapse (n=3). Expression of protein was measured by western blot from formalin embedded samples. (**C**) Venn Diagram showing the overlap in FOXA1 chromatin interactions (ChIP-sequencing) between MCF-7 and BT474 cells. (**D**) Box-plot indicating the binding intensity of FOXA1 at chromatin regions identified in MCF-7 and BT474 cells. The FOXA1 binding of MCF-7 cells over-expressing HER2 were also analyzed at the same chromatin regions identified at section. (E) Average signal intensities of FOXA1 binding sites in control treated cells and in cells treated with HER2 inhibitor (Trastuzumab).

### FOXA1 mediates the proliferation driven by HER2 signaling pathway

Next we aimed to determine the functional significance of FOXA1 binding regulated by HER2. For that, we first assessed the effects of FOXA1 on global gene expression in MCF-7 and BT474 breast cancer cell lines. Since FOXA1 mediates estrogen-induced transcriptional activity (2) and estrogen-ER alters the genomic distribution of FOXA1 (28), we analyzed the gene transcripts regulated by FOXA1 without estrogen treatment. As control, we also included the MDA-MB-453 cell line, which is negative for the expression of ER but still preserves the expression of FOXA1 and intermediate levels of HER2 compared to MCF-7 and BT474 cell lines (Supplementary Figure S3A). Hence, we transfected hormone-deprived breast cancer cells with siControl or siFOXA1, performed gene expression analysis and identified genes and pathways that were FOXA1 regulated (Figure 2A and Supplementary Figure S3B-D). Specific silencing of FOXA1 affected the transcription of genes enriched in cell proliferation in all the breast cancer cell lines tested independently of ER or HER2 status (Figure 2B and Supplementary Figure S3C). Subsequently, we analyzed the distribution of FOXA1 binding sites with regards to FOXA1 regulated genes. We integrated FOXA1 binding regulated by HER2 signaling (from Figure 1C) with FOXA1 regulated genes in a window of ±20Kb from their transcription start sites (TSS). Around 43% of FOXA1 regulated genes in BT474 contained FOXA1 sites regulated by HER2 (Supplementary Figure S4A). Around 34% of FOXA1 regulated genes in MCF-7 also comprised FOXA1 sites (Supplementary Figure S4A). Then, we determined how the inhibition of HER2 impacted the binding of FOXA1 within FOXA1 regulated genes in MCF-7 cells. We detected that the FOXA1 signal was almost abridged at the promoter and the gene body regions of FOXA1 regulated genes upon HER2 inhibition (Figure 2C), suggesting that FOXA1 function may be also regulated by HER2 signaling pathway in ER positive tumors and also displaying low levels of the tyrosine kinase receptor. Then, we investigated how the HER2-regulated FOXA1 binding to the chromatin relates to its cell-specific functions by performing ingenuity pathway analysis of FOXA1 dependent genes with FOXA1 sites controlled by HER2. The top signaling pathways associated with FOXA1 regulated genes included cell cycle regulation, estrogen-mediated S-phase entry and breast cancer pathogenesis (Figure 2D).

**Figure 2.**
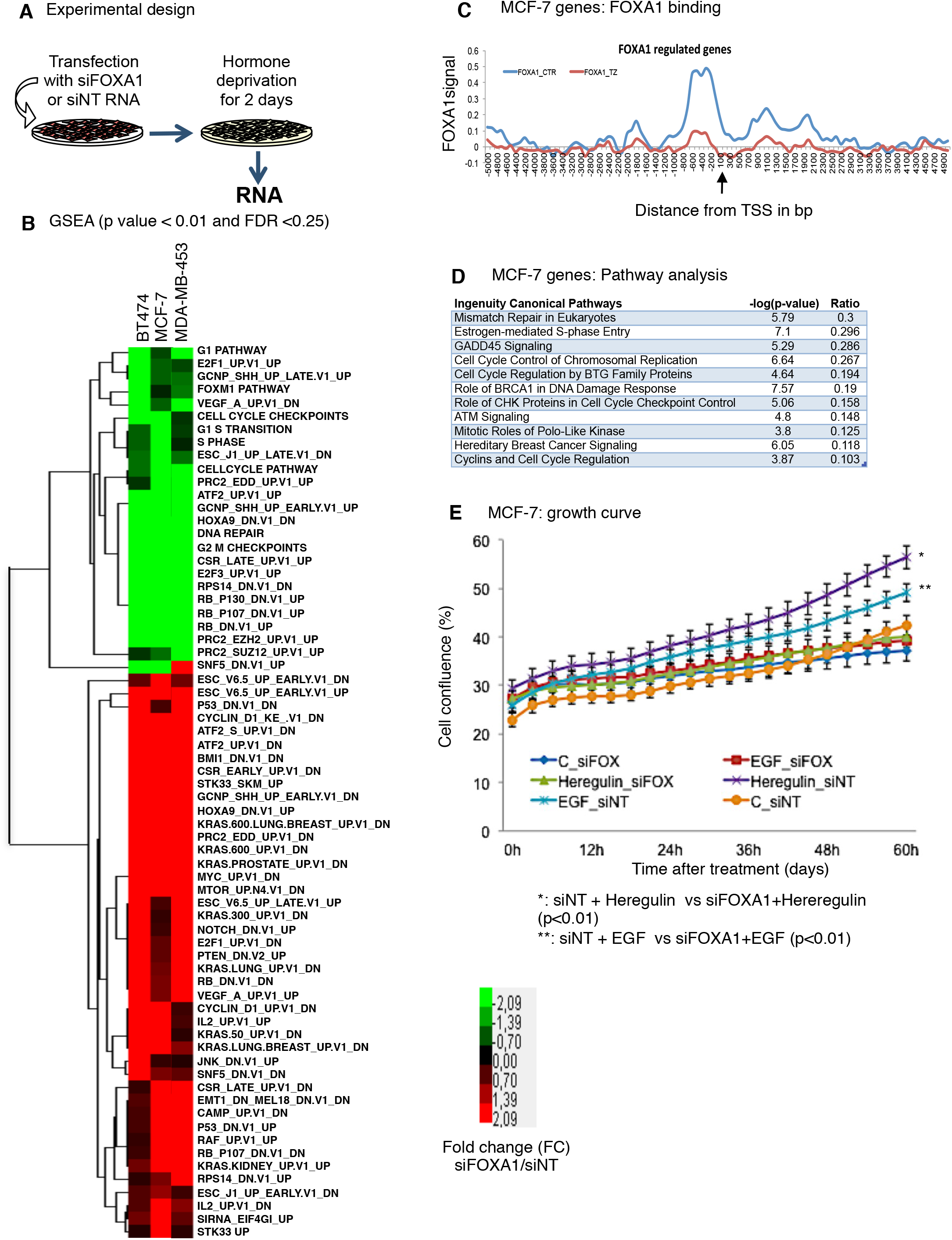
FOXA1 mediates proliferation triggered by HER signaling in ER positive breast cancer cells. (**A**) Overview of the workflow. Hormone deprived breast cancer cells were transfected with siFOXA1 or siControl RNA for 48 h and total RNA was collected for microarray experiments. (**B**) Heatmap of biological processes significantly altered after knocking down FOXA1 in three cell lines showing the overview of the effect on a selection of gene sets that were run in GSEA (selection; p value ≤ 0.01, FDR ≤ 0.25). Green represents downregulation, red upregulation and black no effect. (**C**) The relative difference in FOXA1 signal of promoter regions of genes regulated by FOXA1 in MCF-7 cells. Cells were treated with Control or with HER2 signaling pathway inhibitor Trastuzumab. (**D**) Ingenuity Pathway Analysis of FOXA1 dependent genes containing FOXA1 sites regulated by HER2 signaling pathway in MCF-7 cells. (**E)** MCF-7 cells were transfected with siControl or siFOXA1 and treated with vehicle, EGF or Heregulin. Total cell growth was assessed. The data are the mean of independent replicates ± s.d.

Previously, it was reported that EGFR/HER2 signaling was responsible for ER activation (29). Moreover, we previously demonstrated that FOXA1 is key determinant of ER function in breast cancer (30). Hence, we hypothesized that FOXA1 might be mediating the signals of HER2. Accordingly, we analyzed the dependence of FOXA1 in the growth of ER positive cells treated with specific growth factors for the activation of EGFR (EGF) or HER3 (Heregulin). We assessed the growth in hormone sensitive (MCF-7) and hormone resistant breast cancer cell lines (TAMR (31) and BT474 (32)). EGF and Heregulin induced cell growth in cells expressing FOXA1, whereas the silencing of FOXA1 abrogated their stimulatory effect (Figure 2E and Supplementary Figure S4B and C). These results support that FOXA1 is instrumental for HER2-mediated cell proliferation and leads to a hormone-resistant context.

### FOXA1 expression in HER2-high breast cancers is associated with poor prognosis and hormone-resistance

To investigate if the FOXA1 regulated genes were functionally relevant in breast cancer progression, we analyzed how many of those genes were represented in a gene expression predictor list of poor prognosis for ER positive tumors (15). This signature consisted of two groups of genes; those with relatively high expression and those with relatively low expression. Accordingly, we performed the analysis in one HER2-low (MCF-7) and two HER2-high/moderate cell lines (BT474 and MDA-MB-453, respectively). We identified that more than 50% of the genes predicting poor prognosis were FOXA1 regulated in any of the cell lines investigated. Interestingly, additional FOXA1-regulated genes were identified in the HER2-high/moderate groups (Figure 3A and Figure Supplementary S5). We also sought to determine if the FOXA1 regulated genes associated with poor prognosis could be differentially expressed in patients with relapse to anti-ER therapies (from Figure 1A). We analyzed a subset of these genes and the results showed that expression of the vast majority of these FOXA1 regulated genes with high expression was increased in metastases compared to primary tumors. Moreover, the expression of FOXA1 regulated genes with low expression was decreased in metastases compared to primary tumors (Figure 3B). These results, together with the data showing increased HER2 expression in metastases from patients that relapsed on endocrine therapy, raised the question whether FOXA1 expression in a HER2-enriched environment could be contributing to a more invasive phenotype. Therefore, we conducted a Gene Set Enrichment Analysis (GSEA) on a signature of genes associated with a metastatic signature in breast cancer (33). The analysis demonstrated that the genes up regulated by FOXA1 in HER2-high cells were significantly enriched in the metastatic signature (Figure Supplementary S6A-C). Next, we decided to test the migration ability of HER2-MDA-MB-453 cells overexpressing FOXA1 and also stimulated with Heregulin or EGF. The overexpression of FOXA1 alone did not influence the migration of cells compared to non-overexpressing cells, but Heregulin treatment induced more migration compared to EGF and it was even larger in FOXA1 overexpressing cells (Figure 3C). Moreover, the migration triggered by Heregulin was abridged when FOXA1 was depleted (Figure Supplementary S6D). Altogether these results support that FOXA1 contributes to control the expression of genes associated with poor prognosis in tumors with enhanced HER2/HER3 signaling.

**Figure 3.**
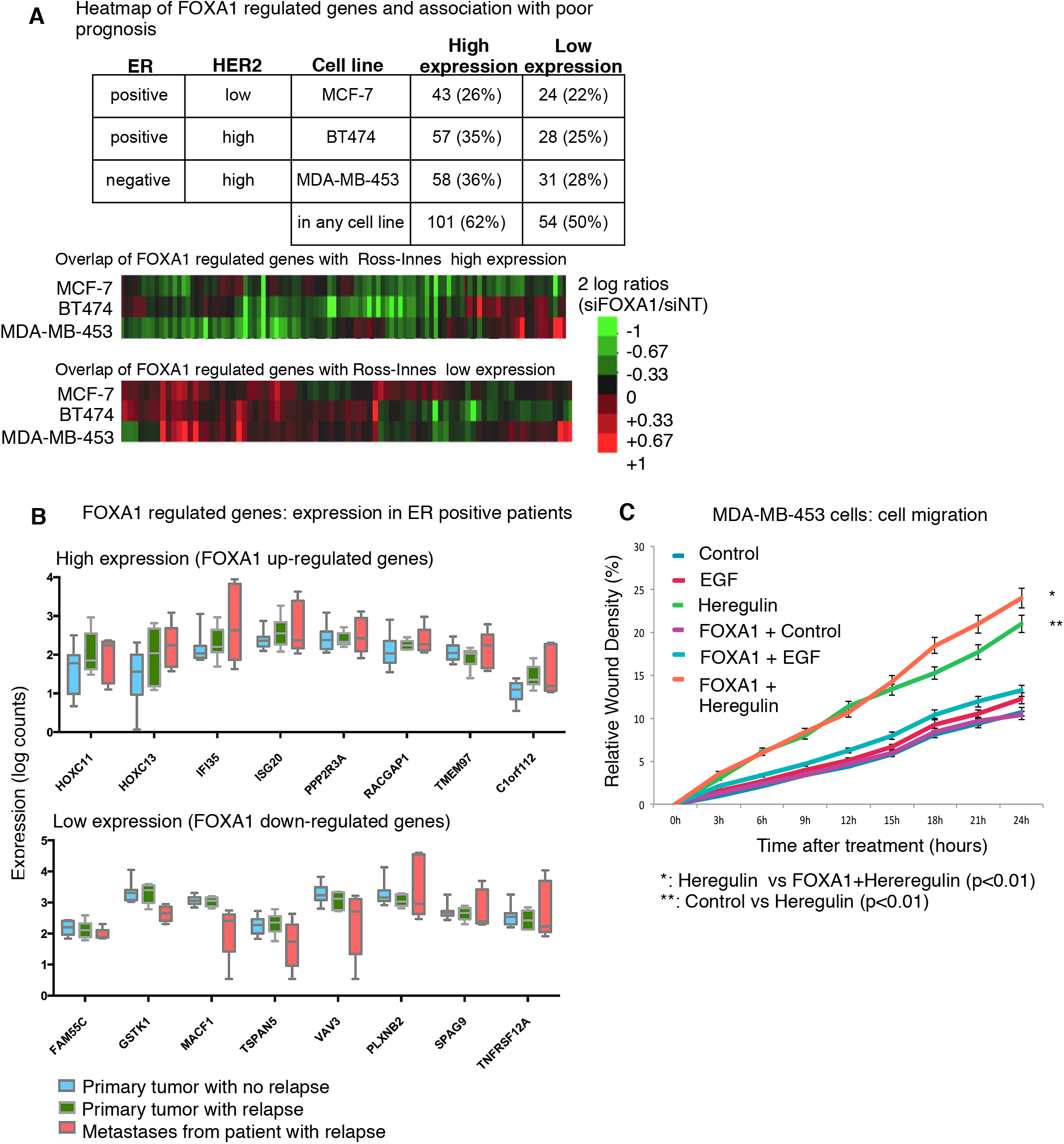
FOXA1 triggers transcription of poor prognosis genes in HER2-high cells. (**A**) The percentage of significantly (P value <0.05) differentially expressed genes in Ross-Innes poor prognosis gene signatures (15) (with positive or negative expression) after knocking down FOXA1 in breast cancer cells. The heat-maps represent 2 log ratios of siFOXA1/siControl of differentially expressed genes upon FOXA1 depletion in MCF7 (HER2-low), BT474 and MDA-MB-453 (HER2-high) cell lines. Green represents down-regulation, black represents no change, and red represents up-regulation as indicated at the bar legend. (**B**) Expression analysis of FOXA1 regulated genes associated with poor prognosis in groups of ER positive patients from primary tumors from patients without relapse (n=10) and paired samples of metastases (n=5) and primary tumors from patients with relapse (n=5). Expression of mRNA was measured by using Nanostring techonology from formalin embedded samples. (**C**) Effect of FOXA1 and Heregulin on migration of breast cancer cells. MDA-MB-453 cells stably expressing a *Doxycycline-inducible* FOXA1 gene were treated with vehicle (FOXA1+Control), EGF (FOXA1+EGF) or Heregulin for 24 hours. Non-inducible doxycycline cells and treated with vehicle (Control), EGF or Heregulin for 24 hours were used as control. The data are the mean of three independent replicates ± s.d. Lower panel shows FOXA1 protein levels with western blotting (HA antibody) and actin as loading control.

### HER3 confers ER independent growth and FOXA1 mediates this effect

Our results show that HER2 expression is increased in metastases from patients with relapse on anti-ER therapies (from Figure 1A,B) and that HER2/HER3 signaling switches the role of FOXA1 to be a less likely regulator of ER (from Figure 3C). These results suggests that by enhancing HER2/HER3 signaling in luminal-like breast tumor cells, FOXA1 might drive tumor growth in an ER independent manner, which might explain the lack of response to anti-ER therapies. To test this hypothesis, we used a previously established luminal-like breast cancer Patient Derived Xenograft (PDX) mouse model (25), in which the tumor growth is still dependent on estrogen after several passages *in vivo*. The mice were treated with vehicle, Fulvestrant, Trastuzumab, EGF, Heregulin and combinations of these. Tumor size was measured at regular intervals during treatment in all animals and tumor growth was analyzed across time (Figure Supplementary S7 and Supplementary table 2). Treatment with Fulvestrant led to cessation of tumor growth compared with the untreated animals in which tumor volume approximately doubled in 20 days (Figure 4A). Treatment with Heregulin led to increased tumor growth, while EGF treatment reduced tumor growth slightly compared to control mice. Combining Fulvestrant with either of these two growth factors revealed a striking difference; while EGF combined with Fulvestrant decreased tumor growth to a level similar to those receiving Fulvestrant monotherapy, Heregulin in combination with Fulvestrant resulted in a growth rate almost at the level of control mice. We corroborated these experiments in breast cancer cell lines (Figure Supplementary S8A). Importantly, the tumor growth triggered by HER2/HER3 in ER depleted tumors was abridged when animals were treated with HER2 inhibitor (Figure 4A). Next, we performed FOXA1 ChIP-seq in the PDX tumors and compared FOXA1 binding between the different treatment groups (Figure 4B). The overlap among these groups revealed that FOXA1 binding was increased in tumors from animals treated with Fulvestrant. Interestingly, the increase of FOXA1 binding was even larger when animals were treated with Fulvestrant and Heregulin (Figure 4C and D) and its increased binding was found to be enriched towards gene transcripts associated with poor prognosis in breast cancer (Figure Supplementary S8C). Moreover, we analyzed the genomic distribution of FOXA1 sites from Figure 4B and compared among the treatments. The results showed that FOXA1 had an increased binding at gene body with the treatment of Heregulin compared to control (Figure Supplementary S8D). Importantly, these data are in agreement to the results obtained in cell lines, where the increased levels of HER2 resulted in a reprograming of FOXA1 binding. Next, we determined the effects of ER depletion by Fulvestrant treatment on gene expression of some of these genes. Under ER depleted conditions, FOXA1 was still expressed (Figure Supplementary S8B), but treatment with Heregulin induced expression of genes associated with poor prognosis, whereas EGF rarely showed such effect (Figure 4E). These results demonstrated that HER2/HER3 activation in ER positive tumors might overcome the tumor growth arrest in patients treated with anti-ER drugs. Importantly, FOXA1 binding to the chromatin in these tumors was regulated by the HER2/HER3 signaling pathway (Figure 4C), which supports that FOXA1 mediates the proliferative signals of HER2/HER3 even when ER is not expressed. Consistent with these results, analysis of gene expression in breast tumors from the METABRIC cohort, revealed significant (P<0.001) correlations between the expression of FOXA1 and HER3 in luminal and HER2-enriched subtypes. Notably, the correlation was higher in the HER2-enriched subtype when compared with the luminal subtypes (Figure Supplementary S9).

**Figure 4.**
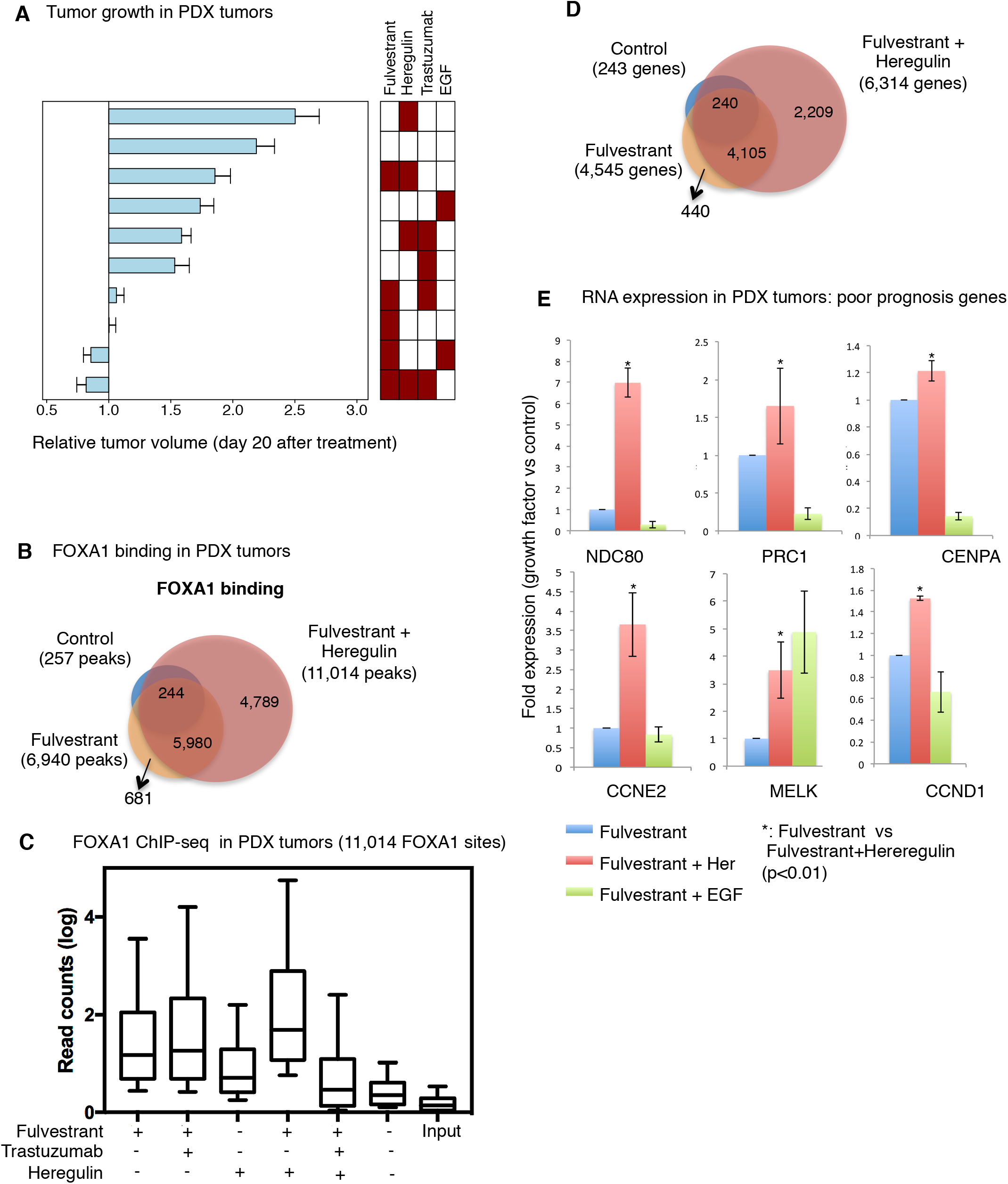
Enhanced activity of HER3 signaling pathway overcomes ER inhibition in a PDX model of a luminal-like breast tumor. (**A**) Change in tumor volume across the first 20 days after pump insertion and initiation of treatment. The bars are sorted according to relative change in tumor volume over time and the heatmap illustrates the corresponding treatment combinations. (**B**) Venn Diagram showing the overlap in FOXA1 chromatin interactions (ChIP-sequencing) at PDX tumors from control animals and treated with Fulvestrant or Fulveltrant and heregulin. (**C**) Box-plot indicating the binding of FOXA1 to the chromatin related to heregulin induced sites in PDX tumors with different treatments. Wilcoxon rank-sum test was used to test any statistical difference between samples. (**D**) Venn Diagram showing the overlap of genes with FOXA1 binding identified in panel c. (**E**) Real-time PCR of genes associated with poor prognosis. RNA from PDX tumors treated with fulvestrant, fulvestrant and EGF or fulvestrant and Heregulin was collected. The data are represented as the mean of independent replicates ± s.d.

### HER2/HER3 facilitates chromatin accessibility of FOXA1 binding regions with reduced binding of ER and EP300

Altogether, the results of this study support that breast tumors and cell lines initially sensitive to anti-estrogen drugs may overcome this inhibition by enhancing HER2/HER3, either by increasing the levels of these receptors or their activating ligands. Therefore, we next aimed to gain insight into the mechanisms by which FOXA1 might confer the ability of ER positive tumors to grow independently of ER. Given the differences observed in the FOXA1 chromatin binding between HER2-high vs. HER2-low cell lines, we hypothesized that in the former we could be observing an additional effect which could ultimately explain the ER independent role of FOXA1. Consequently, we first determined the frequency of recognition motifs for ER (ERE) and FOXA1 (FKH) within the differential FOXA1 binding events detected in HER2-low (MCF-7) and HER2-high (BT474) breast cancer cells (from Figure 1B). The FKH motifs were highly represented independently of FOXA1 binding regions identified in MCF-7 or BT474 cells. By contrast, ERE motifs were unrepresented in FOXA1 unique regions identified at BT474 cells (Figure 5A left panel), which suggested that high HER2 signaling influenced FOXA1 to pioneer genomic regions with reduced ER binding and probably to ER co-regulators. Then, we determined the binding of ER and of the ER co-regulator EP300 towards these FOXA1 regions. The results revealed that BT474 unique regions were poorly enriched of ER and EP300 (Figure 5A right panel), which confirmed our hypothesis. We also investigated the frequency of motifs for other pioneer factors described previously to play a substantial role in breast cancer, such as AP2γ and PBX1 (34). In relation to AP2γ, we identified significant motif enrichment exclusively at FOXA1 sites from BT474 cells (Figure Supplementary S10A). Interestingly, the pioneer function of AP2γ has been reported to be dependent of FOXA1 (35), which suggests that AP2γ might cooperate with FOXA1 in the control of chromatin accessibility regulated by HER2. PBX1 was not significantly enriched in any of the FOXA1 binding regions (Figure Supplementary 10A). Despite PBX1 playing a FOXA1 independent role (36) our results suggest that HER2 might not impact the PBX1 function at FOXA1 binding regions.

**Figure 5.**
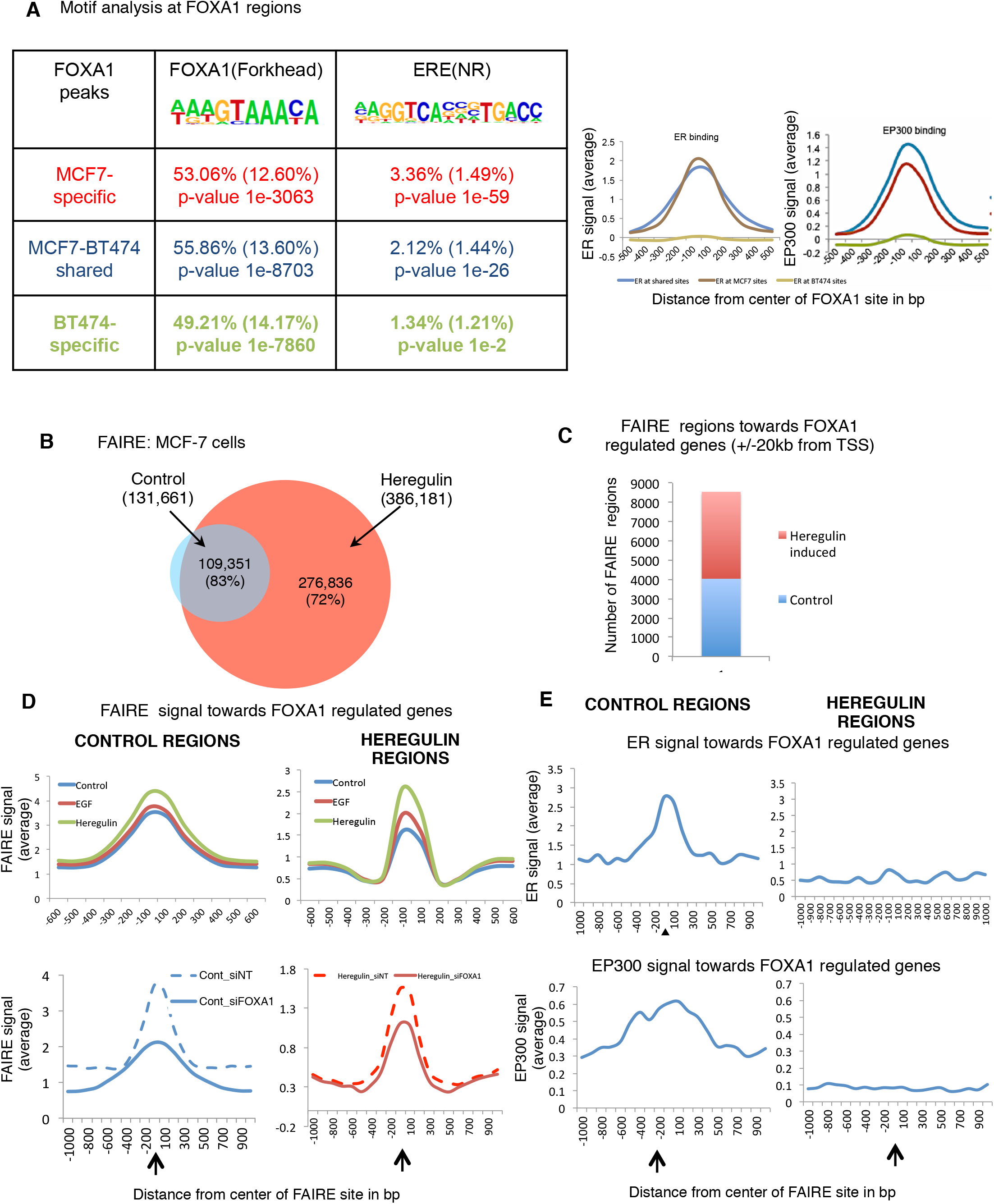
Differential binding of FOXA1 to chromatin is influenced by HER2 expression. (**A**) Determination of the sequence recognized by FOXA1 (FKH) or ER (ERE) within its cell type-specific binding sites. The sequences of FKH and ERE motifs analyzed are shown. The relative difference in ER or EP300 signal within the FOXA1 cell type-specific binding sites is shown. (**B**) Genome-wide FAIRE was performed in MCF-7 cells treated with vehicle (Control) or Heregulin for 1h. Diagram shows overlap in FAIRE regions between vehicle-treated and Heregulin-treated cells. (**C**) Number of vehicle or Heregulin FAIRE regions enriched towards FOXA1 regulated genes (± 20kb from TSS). (**D**) The different categories of FAIRE regions (control, EGF and Heregulin induced) enriched towards FOXA1 regulated genes were assessed. The upper panel represents the average of FAIRE signal in cells treated with vehicle or Heregulin. Lower panel shows the average FAIRE signal in cells transfected with siControl or siFOXA1. (**E**) The upper panel indicates the average binding of ER signal from a previous publication (2) and the lower panel for EP300 for each of the FAIRE categories.

FOXA1 can mimic linker histones and bind directly to compacted chromatin (37), therefore exposing ERE motifs and allowing ER-DNA interaction (2). Hence, we investigated the global impact of HER2/HER3 signaling activation on the pioneering function of FOXA1. Since we have observed that FOXA1 binding is reprogramed in both ER positive cell lines and tumors when HER2/3 signaling is triggered, we investigated the pioneering function of FOXA1 in MCF-7 stimulated with Heregulin. We performed formaldehyde-assisted isolation of regulatory elements (FAIRE) coupled with high-throughput sequencing to identify euchromatic regions of the genome (38). Cells were transfected with siControl or siFOXA1 and then control-treated or treated with Heregulin. In control-treated cells, we found 131,661 FAIRE regions and in Heregulin-treated cells we found 386,181 FAIRE regions. Around 83% of the control-treated FAIRE regions (Figure 5B) were also found in the Heregulin-treated group. Heregulin treatment induced novel FAIRE regions and they represented the 72% of the total regions induced by the growth factor (Figure 5B). We also examined the number of FAIRE regions towards FOXA1 regulated genes. Heregulin induced more euchromatic regions compared to control (Figure 5C). Next, we examined the FAIRE signal triggered by EGF at control and Heregulin-specific regions and compared with Heregulin. The growth factor EGF was able to open the chromatin at control regions but its effect on Heregulin-specific regions was not substantially significant (Figure 5D top panel). Moreover, in cells depleted of FOXA1 we observed substantial decreases in the opening of the chromatin at control FAIRE regions. Interestingly, the FAIRE regions selectively induced by Heregulin were partially regulated by FOXA1, as a subset (36%) of these regions was substantially affected by FOXA1 depletion (Figure 5D bottom panel and Figure Supplementary S10B). Importantly, HER3 triggered signal stimulated FOXA1 binding to chromatin (Figure Supplementary S10C and D), which suggests that HER3 provides a more euchromatin state in a FOXA1-dependent manner. Next, we analyzed the binding of ER and the histone-modified cofactor EP300 within the identified FAIRE regions (Figure 5E). We observed global ER and EP300 binding at control-regions. By contrary, ER and EP300 binding were abridged in Heregulin-induced regions. These results support that the binding of ER and EP300 to HER3-induced regions is less likely to occur, whereas their binding to control regions is not impeded.

### HER2/HER3 confers ER independent cell growth by inhibiting FOXA1 acetylation

Previously, it was described that ER function is dependent of the intrinsic acetyl transferase activity of EP300 (39). Interestingly, EP300 is able to directly acetylate FOXA1 *in vitro*, and EP300 driven acetylation prevents FOXA1 DNA binding, but does not affect the protein when already bound to DNA (40). These reports together with our results made us to postulate that the enzyme EP300 might retain FOXA1 at ER binding regions to mediate ER function. By contrast, the increased signaling of HER2/3 inhibits the acetylation of FOXA1 to enable the binding of the transcription factor to additional regions non-enriched of ER. To test our hypothesis, we first analyzed the acetylation of FOXA1 in ER positive (MCF-7) and ER negative (MDA-MB-453) breast cancer cell lines. We performed immunoprecipitation of FOXA1 followed by western blot of FOXA1 and acetyl lysine modification. FOXA1 was only acetylated in ER positive cell line (Figure 6A) and this acetylation was dependent of EP300 (Figure Supplementary S11A). Moreover, we investigated how FOXA1 acetylation was influenced by HER2/HER3 signaling. First, we treated MCF-7 cells with Heregulin and performed FOXA1 immunoprecipitation. The results revealed that HER2/3 activation prevented FOXA1 acetylation in MCF-7 cells (Figure 6B). Then, we investigated how that HER2 inhibition impacted the acetylation of FOXA1 in ER positive breast cancer cell lines with high HER2 levels (BT474 and MCF7-HER2). The inhibition of HER2 increased the acetylation of FOXA1 in these cell lines (Figure Supplementary S11B and C). Altogether, these results supported that HER2 and ER played opposite roles in the regulation of FOXA1 acetylation.

**Figure 6.**
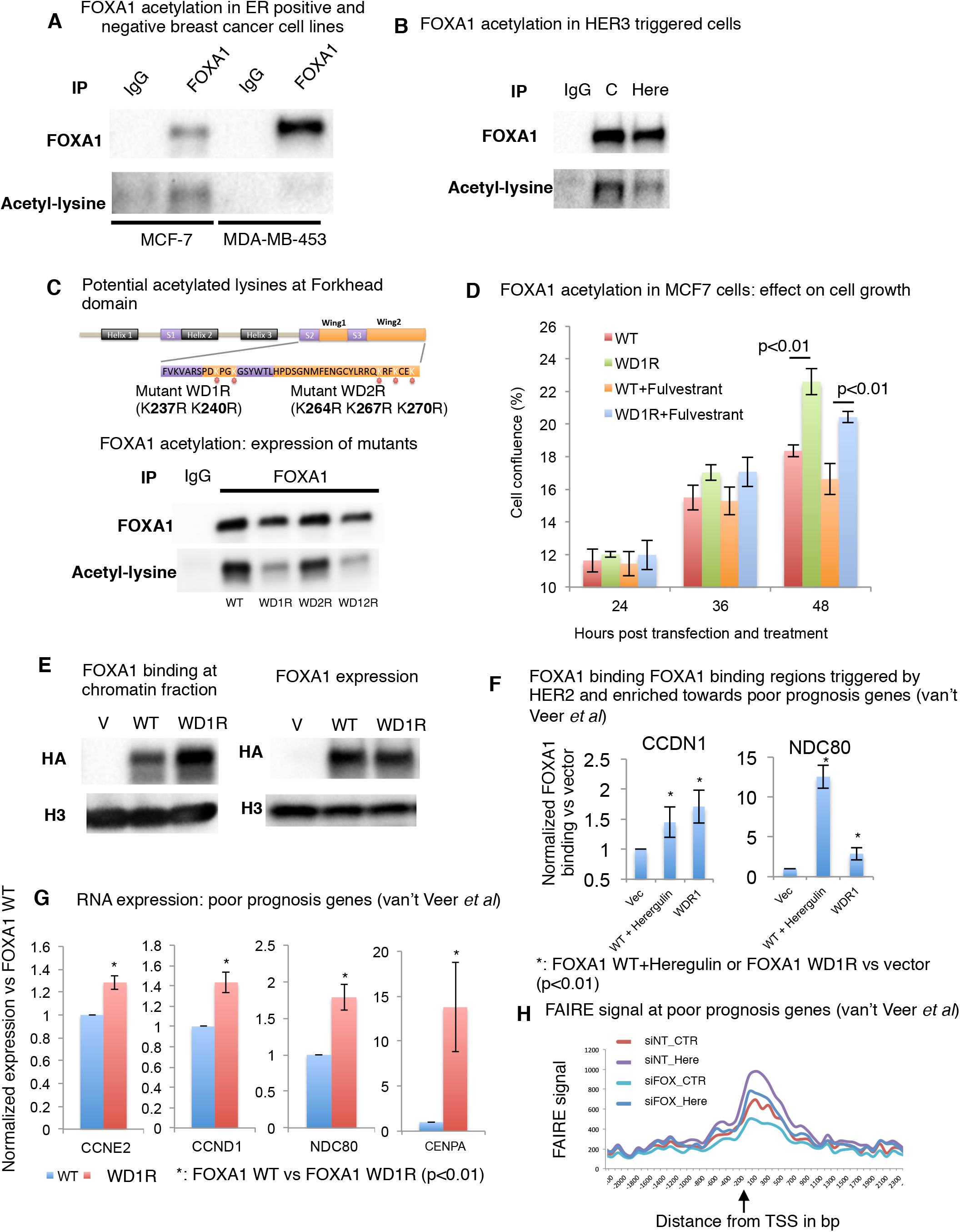
FOXA1 deacetylation correlates with ER independent function. (**A**) FOXA1 acetylation from immunoprecipitated FOXA1 protein complex in MCF-7 and MDA-MB-453 cell lines. (**B**) FOXA1 acetylation from immunoprecipitated FOXA1 protein complex in MCF-7 control treated and treated with heregulin. (**C**) Potential acetylated lysine aminoacids identified at Forkhead domain of FOXA1 (top panel). FOXA1 acetylation from immunoprecipitated FOXA1 protein complex in MCF-7 cells transfected with wild type (WT), lysines mutated into arginines at Wing Domain 1 (WD1R), lysines mutated arginines at Wing Domain 2 (WD2R) and double mutant (WD12R) (bottom panel). (**D**) MCF-7 cells were transfected with vectors expressing FOXA1 WT or FOXA1 mutant WD1R and treated with fulvestrant. The cell growth was measured at different time points. (**E, F**) The global binding of FOXA1 to the chromatin in MCF-7 cells transfected with WT and WD1R FOXA1 mutant was tested. In E, is shown the global FOXA1 binding at chromatin fraction (by western blot). In F, is shown the FOXA1 binding at chromatin binding regions identified at BT474 cells (by ChIP-PCR) in cells transfected with vector, transfected with FOXA1 WT plus Heregulin and transfected with FOXA1 mutant (WD1R). (**G**) Real-time PCR of genes associated with poor prognosis. RNA from MCF-7 cells transfected with WT and WD1R FOXA1 mutant was tested. The data are represented as the mean of independent replicates ± s.d. (**H**) The relative FAIRE signal in control treated cells or treated with heregulin towards genes associated with poor prognosis (van’t Veer *et al*). Red: control treated, purple: Heregulin treated. Light blue: control cells treated with siFOXA1, blue: Heregulin treated cells treated with siFOXA1.

Previously, it has been reported that FOXA1 contains five acetylation sites identified at the wings of the fork-head domain, two at the wing 1 (WD1) and three at the wing 2 (WD2). Interestingly, FOXA1 mutations at the same domain were recently described and these mutations were associated with an increased FOXA1 function (40). We hypothesized that FOXA1 deacetylated might mimic the effect of the mutants identified in breast cancer patients and therefore confer insensitivity to endocrine therapy by inducing tumor growth independently of ER. Accordingly, we first created FOXA1 mutants defective for acetylation at WD1 (WDR1) or WD2 (WDR2) and tested their ability to be acetylated compared to wild type FOXA1 (WT) transfected cells. The mutant for both domains was also created (WDR12) (Figure 6C top panel). Both FOXA1 WT and WDR2 mutant were acetylated but the acetylation of WDR1 mutant was dampened (Figure 6C bottom panel), which supports that WD1 domain is acetylated *in vivo*. Next, we investigated the ability of ER positive cells to growth when they ectopically expressed a FOXA1 mutant defective in acetylation. Cells transfected with FOXA1 mutant increased the cell growth compared to cells transfected with FOXA1 WT. Moreover, the cells transfected with FOXA1 mutant rescued the cell growth arrest in cells depleted of ER (Figure 6D). We also investigated how the mutation deficient in acetylation impacted the binding of FOXA1. We observed a substantial increase of FOXA1 binding to the chromatin for the mutant when it was compared with FOXA1 WT (Figure 6E). Moreover, the mutation increased the binding of FOXA1 at chromatin regions triggered by HER2/3 and with poor ER binding of genes associated with poor prognosis (Figure 6F). Importantly, the expression of genes associated with poor prognosis was also increased in cells transfected with the FOXA1 mutant compared to cells transfected with WT FOXA1 (Figure 6G). Altogether, the results of this study suggest that FOXA1 acetylated by EP300 might retain FOXA1 at chromatin regions where ER also binds to enable ER function. However, when HER2/3 is triggered, FOXA1 acetylation is inhibited and therefore it is able to bind to additional chromatin regions less likely enriched of ER and EP300 to facilitate the transcription of genes associated with poor prognosis. Consistent to this idea, a significant increase of the chromatin accessibility at TSS of genes associated with poor prognosis in cells treated with Heregulin was detected. Importantly, this increase of chromatin accessibility was mediated by FOXA1 (Figure 6H).

In summary, our study demonstrates that hormone-resistant patients with increased signaling of HER2/3, FOXA1 binds to chromatin regions poorly enriched of ER to control the transcription of genes associated with poor prognosis. By doing so, FOXA1 leads to an ER-independent growth and tumors become insensitive to anti-ER drugs.

## Discussion

The results of this study expand the role of HER2 beyond its canonical direct action on ER as previously reported (41). We demonstrate that luminal-like breast tumors acquire the property to grow in an ER independent manner when HER2/HER3 signaling pathway is enhanced. Our results also establish that FOXA1 mediates the HER2/HER3 signaling in a hormone-resistant context. Moreover, we now demonstrate *in vivo* that enhanced activity of HER3 overcomes the inhibitory action of Fulvestrant, by conferring the ability of ER positive tumors to grow in an ER-independent fashion. Preclinical (42) and clinical (43) studies established that the combination of anti-ER with anti-HER2 therapies delays the development of resistance. Our *in vivo* experiments demonstrate that ER inhibition prevents the tumor growth triggered by EGFR/HER2, which support that double targeted therapy (ER and HER2) might inhibit ER dependent growth. However, HER3 mediated signaling hinders this effect and is actually one of the key clinical features identified for resistance to anti-HER2 therapies (44). HER3 works preferably through the dimerization with HER2 (45), supporting that HER3/HER2/FOXA1 axis overcomes the inhibition of the EGFR/ER axis. The consequences of HER2/HER3 signaling could potentially be explained by its impact on FOXA1 in the opening of novel chromatin regions with abridged ER/EP300 binding and facilitating the accessibility of TSS associated with poor prognosis. These results are consistent with previous reports, showing an increased risk for disease progression and a decreased overall survival of patients with HER2-high breast tumors (46) and that hormone-resistant breast cancer cells undergo changes in the chromatin landscape (47).

In this study we demonstrate that HER2/3 signaling inhibits the acetylation of FOXA1 and that the FOXA1 mutant defective for acetylation has increased ability to bind to the chromatin, which might be one of the mechanism by which FOXA1 binds to non-ER enriched sites. Therefore, one might postulate that ER/EP300 restricts the binding of FOXA1 and therefore alterations in the function of EP300 might facilitate FOXA1 function independently of ER. The role of EP300 mediating the ER-transcription has been widely described. In fact, EP300 is a component of the ER co-activator complex (48) and its intrinsic histone acetyl transferase activity influences gene expression of ER (49). Moreover, the interaction of ER with chromatin is mediated by FOXA1 (50), which supports our findings that EP300 acetylates FOXA1 as a mechanism that facilitates the transcription of ER-regulated genes. Hence, our results together with previous reports made us hypothesize that FOXA1 acetylated by EP300 might be retained at chromatin regions where ER also binds and then facilitate the transcription of ER target genes.

Ross-Innes *et al*. established that hormone-resistant breast cancers still recruit ER to the chromatin (15). Our results support that ER-DNA interaction is not impeded by EGFR or HER3 signaling, which suggests that either ER/FOXA1 complex or FOXA1 alone contribute to the expression of genes conferring poor prognosis for breast cancer patients. The role of FOXA1 contribution to breast cancer progression independently of ER is unforeseen, because works published to date have linked FOXA1’s function with ER. However, several studies reported that between 7-25% of patients show discordance in ER expression between a primary tumor and distant metastasis (51). We report several lines of evidence supporting an alternative role of FOXA1. First, the expression of genes associated with poor prognosis is FOXA1 dependent. Second, a significant correlation between the expression of FOXA1 and the expression of the most common isoforms of HER3 is found in HER2-high breast cancer subtypes. Third, HER3 triggers FOXA1 binding to the chromatin and ultimately cell proliferation and migration in an ER independent manner.

In our study we report that *in vivo* Trastuzumab treatment inhibits FOXA1 binding to chromatin when it is triggered by HER2/HER3. Importantly, in these experiments the PDX were treated with Heregulin combined with Trastuzumab, which confirms that activation of HER3 is abridged when animals are treated with anti-HER2 therapy. Previously, Trastuzumab has been described not to block the formation of HER2/HER3 heterodimers (52), which might be in opposition to our *in vivo* experiments. One possible explanation for this contradiction might be that Trastuzumab does not inhibit HER2/HER3 dimerization as the mechanism of HER2 inhibition but rather causes HER2 degradation (53). In fact, the treatment of cell lines with Trastuzumab down-regulates the protein levels of HER2, which confirms our hypothesis. Therefore, Trastuzumab might indirectly inhibit HER3 since HER3 only function as a specialized allosteric activator of other HER proteins (54).

The results of our study demonstrate that ER positive tumors may develop the ability to grow independently of ER, which may have implications for targeted therapy. Our data provide insight into tumor heterogeneity, revealing mechanistic insight into what tumors should be targeted with specific HER-targeted therapies and defining whether ER is still functional and still valid as a single drug target. In current clinical practice, hormone-therapy is sequentially used in the treatment at the second and even at the third line of treatment for patients with advanced disease. Nevertheless, the results of these second/third endocrine lines are globally poor, with clinical responses ranging to 0.4% to 13% and with median Progression Free Survival (PFS) of only 2.8 to 4.5 months (55). Furthermore, according to these data, a substantial number of patients (between 20 and 40%) had progressive disease at first tumor assessment, which implies that a substantial proportion of these patients would benefit of better strategies. Prospective randomized studies comparing the combination of anti-HER2 therapies with aromatase inhibitors (AI) have described modest benefits in PFS with respect AI alone (56,57). However, these benefits are improved when a dimerization inhibitor agent such as Pertuzumab is added in addition (Arpino, G. *et al* PERTAIN study, San Antonio Breast Cancer Symposium-SABCS 2016). From these results we can speculate that treatment with drugs targeting HER3 might serve as additional strategies for the treatment of hormone resistant breast cancer. Additional results (Higgins, M. *et al* SABCS 2014) incorporating novel anti-HER3 therapies support our hypothesis. For example, Serintumumab, a novel antibody that targets HER3, has been investigated in a population of ER positive breast cancer patients that had progressed following hormonal therapies. The combination of Serintumumab and Exemestane demonstrated higher efficacy than Exemestane alone (74% reduction in progression-free survival p<0.003), which confirms that the addition of anti-HER3 therapy can restore endocrine sensitivity to Exemestane. Given the complexity of HER2 signaling and the results of our work, we believe that Fulvestrant may be particularly suited for the combination therapy with other novel targeted treatments for patients who relapse to the first line of treatment. However, whether HER3 direct inhibition in combination with Fulvestrant is the most suitable therapeutical approach needs to be addressed. Future studies should include biomarkers that assist the clinicians for the most optimal therapeutical approach for patients who relapse to the first line of endocrine treatment.

## ACKNOWLEDGEMENTS

We thank Dr. Sandra Lopez and Dr. Ian G. Mills for reading of the manuscript. We thank Elena Gonzalez for her technical assistance. The sequencing service was provided by the Norwegian Sequencing Centre (www.sequencing.uio.no), a national technology platform hosted by the University of Oslo and supported by the “Functional Genomics” and “Infrastructure” programs of the Research Council of Norway and the Southeastern Regional Health Authorities. We acknowledge support from University of Oslo, Research Council of Norway and Helse Sør-Øst authority. M.R.K. was supported by NCMM and S.V.S. is supported by Norwegian Cancer Society.

All the *in vitro* experiments were conceived by M.R.K., S.G., K.K.S., M.B., A-L.B-D. and A.H. All the *in vitro* experiments were conducted by M.R.K., S.G., E.F., S.W. The *in vivo* experiments were conceived by J.H.N., H.B., O.E., T.S. and A.H. All the in vivo experiments were conducted by J.H.N., H.B. and H.Z.T. Computational analyses were conducted by M.R.K, B.B., Y.S. and S.N. Manuscript was written by A.H., M.R.K. and S.G. with help from the other authors.

## SUPPLEMENTARY FIGURE LEGENDS

Figure Supplementary S1. Western blot indicating the protein expression of HER2. (**A**) and quantification of protein levels (**B**) in paired samples of metastases (n=3) and primary tumors from patients with relapse (n=3).

Figure Supplementary S2. (**A**) Genomic distributions at each of the FOXA1 sites identified at MCF-7 or BT474 cells. The genomic distribution has been determined by using Homer. (**B**) Venn Diagram showing the overlap in FOXA1 chromatin interactions (ChIP-sequencing) between BT474 and MCF7-HER2 cells (top panel) and between MCF-7 and MCF7-HER2 cells. (**C**) Western blot of FOXA1 in both cell lines tested and treated 48-h with control or Trastuzumab. 1Z-Actin was used as loading control. (**D**) Western blot of HER2 protein levels in MCF-7 cells with different treatments: control, Trastuzumab and Lapatinib., MCF7-HER2 cells were included as positive control. Histone H3 was used as loading control.

Figure Supplementary S3. Complementary information of figure 2. (**A**) HER2, ER and FOXA1 protein levels measured by western blot in MCF7-HER2, MCF-7, TAM-R, MDA-MB-453 and BT474 breast cancer cell lines. (**B**) The number of significant and differentially regulated genes was selected after FOXA1 depletion (False discovery rate 5%). Western blot to confirm the silencing of FOXA1 and loading control (1Z-Actin) are shown. (**C**) Venn diagram illustrating the number of commonly down-regulated genes after FOXA1 knockdown in MCF-7, BT474 and MDA-MB-453 cell lines. (**D**) Ingenuity pathway analysis performed on commonly affected genes (127 genes). Genes related to DNA repair, hereditory breast cancer signaling, Estrogen mediated s-phase entry, cell cycle, purine nucleotide synthesis, aryl hydrocarbon signaling were identified. Most of these pathways were also significantly associated with genes involved in breast cancer pathogenesis. The top signaling pathways associated with FOXA1 regulated genes included cell cycle regulation and estrogen-mediated S-phase entry.

Figure Supplementary S4. Complementary information of figure 2. (**A**) Fraction of genes significantly regulated by FOXA1 and with HER2 regulated FOXA1 site (Blue). (**B**) Western blot showing HER2, phospho-HER2, HER3 and phospho-HER3 protein levels in MCF-7 cells stimulated with EGF (1h), Heregulin (1h) or Heregulin plus Trastuzumab (1h). (**C**) TAMR and BT474 cells were transfected with siControl or siFOXA1 and treated with EGF or Heregulin. Total cell growth was assessed. The data are the mean of independent replicates ± s.d.

Figure Supplementary S5. Unsupervised hierarchical clustering based on the expression of FOXA1 regulated genes in a signature of genes associated with poor prognosis (15). The clustering was made with Pearson correlation. Heat-maps showing differentially expressed both induced (157 genes) and repressed (107 genes) genes after knocking down FOXA1 in MCF7 (HER2-low), BT474 and MDAMB453 (HER2-high) cell lines. The heat-maps contain 2 log ratios of siFOXA1 vs siNTControl. Green represents downregulation, black represents no change, and red represents upregulation. The intensity of the color related to the 2log ratio of up- or down-regulation is indicated by the bar.

Figure Supplementary S6. Gene Set Enrichment Analysis of a metastatic signature (24). (**A**) The list of the 55 poor prognosis prediction genes obtained from the study. (**B**) GSEA graph represents enrichment for vant Veer poor prognosis gene signature in BT474 cells after knocking down FOXA1, the list of genes contributed for core enrichment. (**C**) GSEA graph represents enrichment for vant Veer poor prognosis gene signature in MDA-MB-453 cells after knocking down FOXA1, the list of genes contributed for core enrichment. NES, normalized enrichment score and FDR. The name of shared enriched genes in both cell lines is high lined in red. (**D**) MDA-MB-453 cells were transfected with siControl or siFOXA1 and treated with EGF or Heregulin. Cell migration was assessed. The data are the mean of independent replicates ± s.d at 24h of migration.

Figure Supplementary S7. Relative tumor growth rates under the various treatments. The lines represent growth rate modeled by linear regression and the dots represent individual measurements of relative tumor volume at the indicated day calculated from each tumor measurements. (**A**) Red, Fulvestrant+Heregulin+Herceptin; Orange, Fulvestrant; green, Fulvestrant+Heregulin, grey, control. (**B**). Orange, Fulvestrant; green, Herceptin; grey, control; pink, Heregulin. (**C**) Light blue, Fulvestrant+EGF; orange, Fulvestrant; blue, EGF; grey, control. The bands around each curve correspond to +-1 s.e.

Figure Supplementary S8. Enhanced activity of HER signaling pathway rescues ER inhibition in breast cancer cells. (**A**) Left panel: Western blot confirming reduced levels of ER after treatment of Fulvestrant. Right panel: Cell proliferation of MCF-7 or BT474 with EGF or Heregulin in the presence or absence of fulvestrant. (**B**) Western blot analyses of ER and FOXA1 protein expression in PDX tumors from vehicle-(CTR), fulvestrant-, EGF plus fulvestrant- or Heregulin plus fulvestrant-treated mice. (**C**) Box-plot indicating the binding of FOXA1 to the chromatin related to heregulin induced sites enriched towards genes associated with poor prognosis (van’t Veer et al). in PDX tumors with different treatments. Wilcoxon rank-sum test was used to test any statistical difference between samples. (**D**) Genomic distributions at each of the FOXA1 sites identified at MCF-7 or BT474 cells. The genomic distribution has been determined by using Homer.

Figure Supplementary S9. Complementary information of figure 4. (**A**) Genomic location of the FOXA1 probe and the three ERBB3 probes analyzed for expression in the Metabric project (9,46) (ILMN_173993, ILMN_2397602, ILMN_1751346). Probe ILMN_1751346 recognizes the *ERBB3* mRNA isoforms (UCSC Genes): uc010spb, uc009zok, uc001sjk and uc001sjl. (**B**) Linear regression analysis of FOXA1 and *ERBB3* expression probes in the luminal A, luminal B and HER2-enriched breast cancer subtypes separately. ERBB3 probe ILMN_1751346 showed a higher correlation with FOXA1 expression in the HER2-enriched subtype compared with the luminal subtypes.

Figure Supplementary S10. (**A**) Motif analysis within FOXA1 differentially regions regulated by HER2. PBX1 and AP21⍰ frequency motifs were determined. (**B**) Non-FOXA1 dependent FAIRE regions enriched towards FOXA1 regulated genes (64%). FAIRE signal was measured in MCF-7 cells transfected cells with siFOXA1 or siControl. The average of FAIRE signal for heregulin regions was measured in heregulin treated cells. (**C**) HER2-high (BT474) and HER2-low (MCF-7 cells) were treated with vehicle (non-treatment), EGF or Heregulin for 60min. Western blot of protein at chromatin-enriched fraction was tested for FOXA1. H3 was used as loading control. (**D**) HER2-high (BT474) and HER2-low (MCF-7 cells) were treated with vehicle (non-treatment), or Heregulin for 60min. FOXA1 binding (by ChIP-PCR) was determined at BT474 unique regions.

Figure Supplementary S11. Related to figure 6. (**A**) Stable MCF-7 cells expressing FOXA1-HA were transfected with siControl RNA or siEP300 RNA. The EP300 protein levels were determined by western blot. Actin was used as a loading control (right panel). FOXA1 acetylation from immunoprecipitated FOXA1 protein was determined by western blot (left panel). The total HA-FOXA1 protein levels were also analyzed. (B and C) FOXA1 acetylation from immunoprecipitated FOXA1 protein was determined by western blot in MCF7-HER2 (**B**) and BT474 (**C**) cell lines and compared to cells treated with anti-HER2 drugs Trastuzumab (Her) or Lapatinib (Lap). In both cell lines the HER2 and phosphor-HER2 protein levels were determined by western blot. Actin was used as a loading control (right panel). FOXA1 acetylation from immunoprecipitated FOXA1 protein was determined by western blot (left panel).

