## Supplementary figures and tables for "Breast tumors escape endocrine therapy by ER-independent mechanisms triggered by the coordinated activities of HER2/HER3 and deacetylated FOXA1"

**Figure Supplementary 1**

**A** HER2 protein expression in paired samples of patients

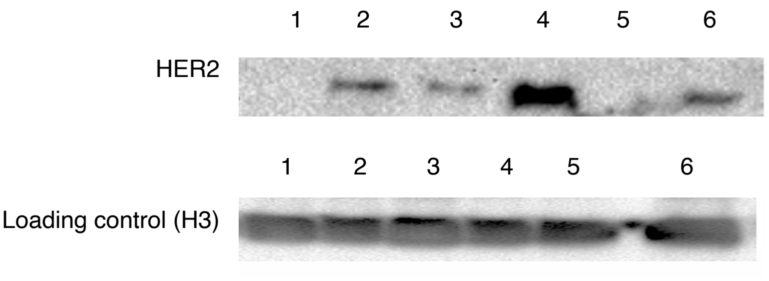


**B** Quantification of HER2 signal vs. loading control

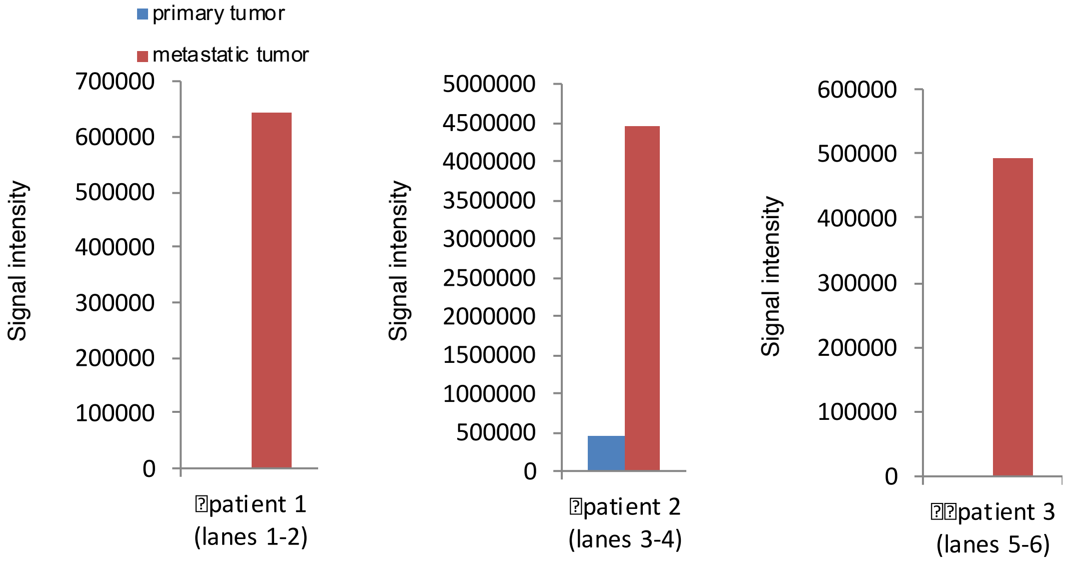

**Figure Supplementary 2**

**A** Genomic distribution of FOXA1 binding sites

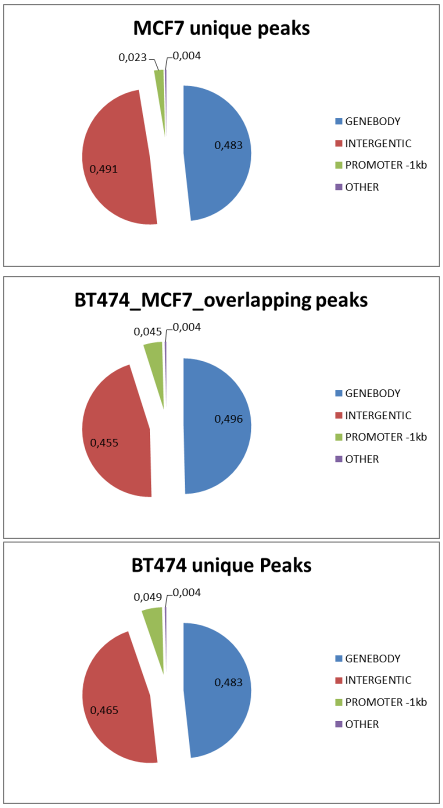


**B** FOXA1 binding in breast cancer cell lines

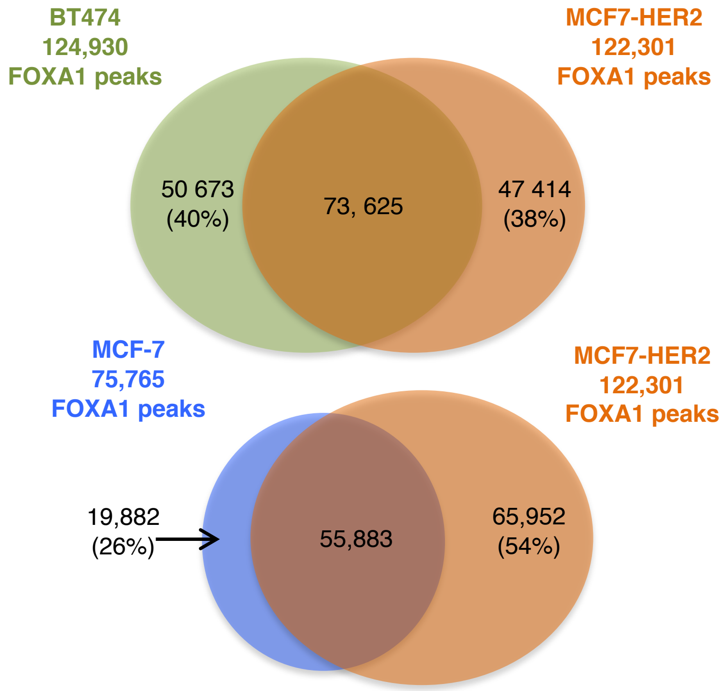


**C** FOXA1 protein upon HER2 inhibition

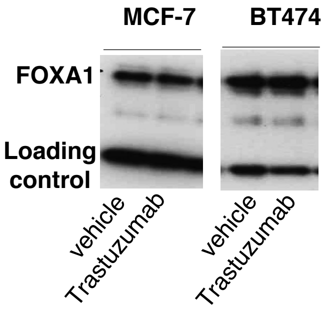


**D** HER2 protein levels in MCF-7 and MCF7-HER2

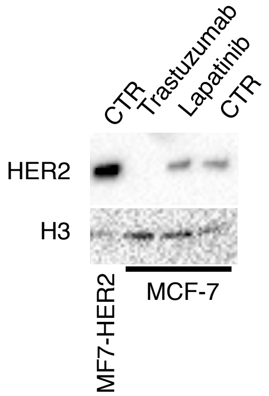

**Figure Supplementary 3**

**A** HER2, ER and FOXA1 protein levels in breast cancer cell lines

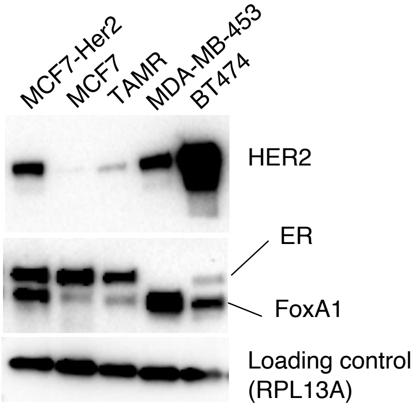


**B** FOXA1 regulated genes (4 replicates)

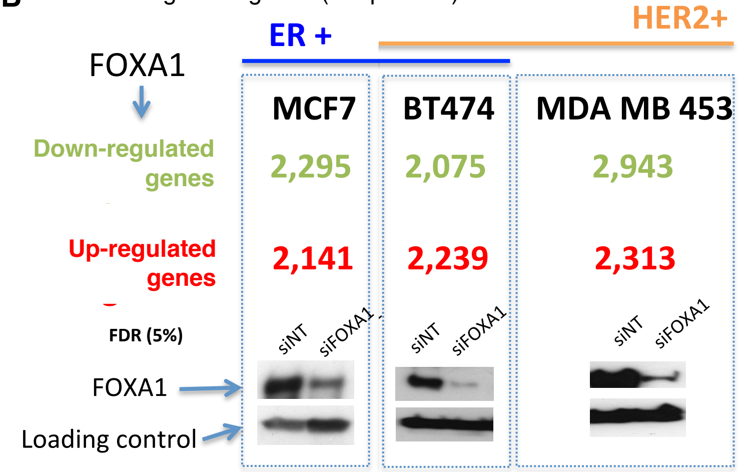


**C** Down-regulated genes in FOXA1 depleted cells

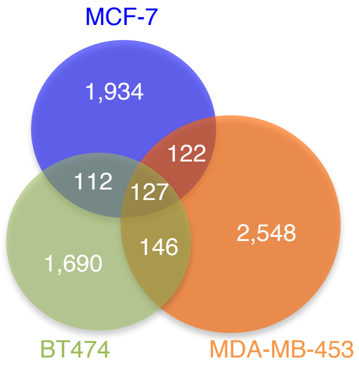


**D** Integrated Pathway Analysis of the common down-regulated genes

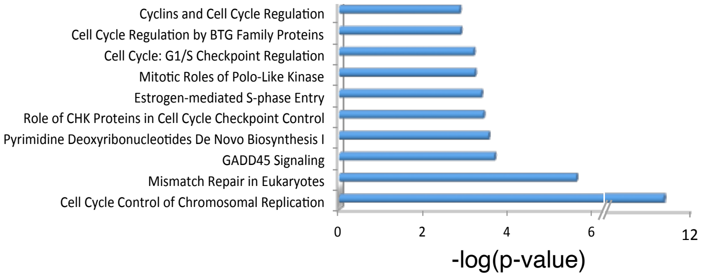

**Supplementary Figure 4**

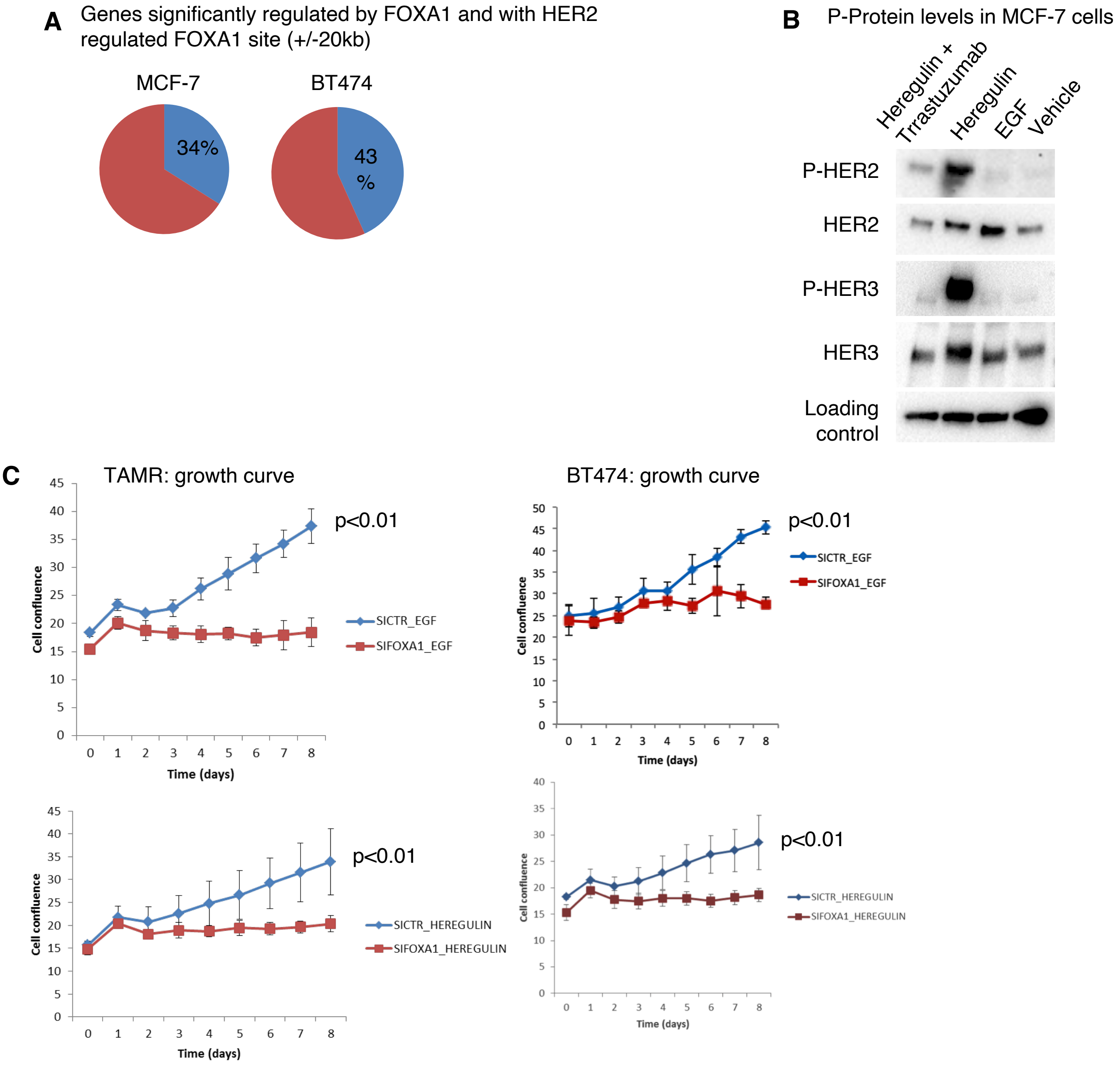

**Figure Supplementary 5**

GSEA of poor prognosis signature (Ross Innes *et al*)

High expression (157 genes)

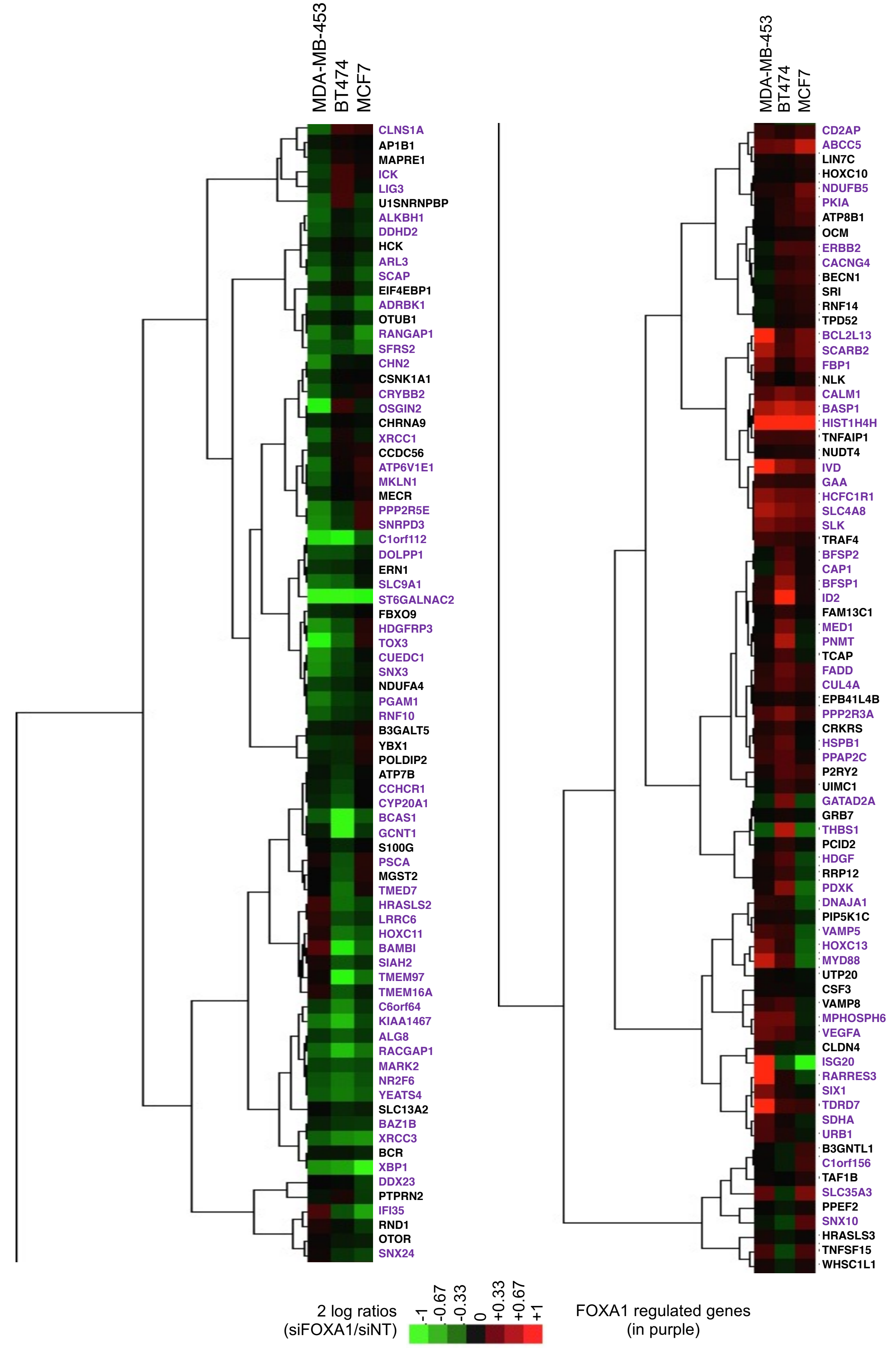

#### Continuation of Figure supplementary 5

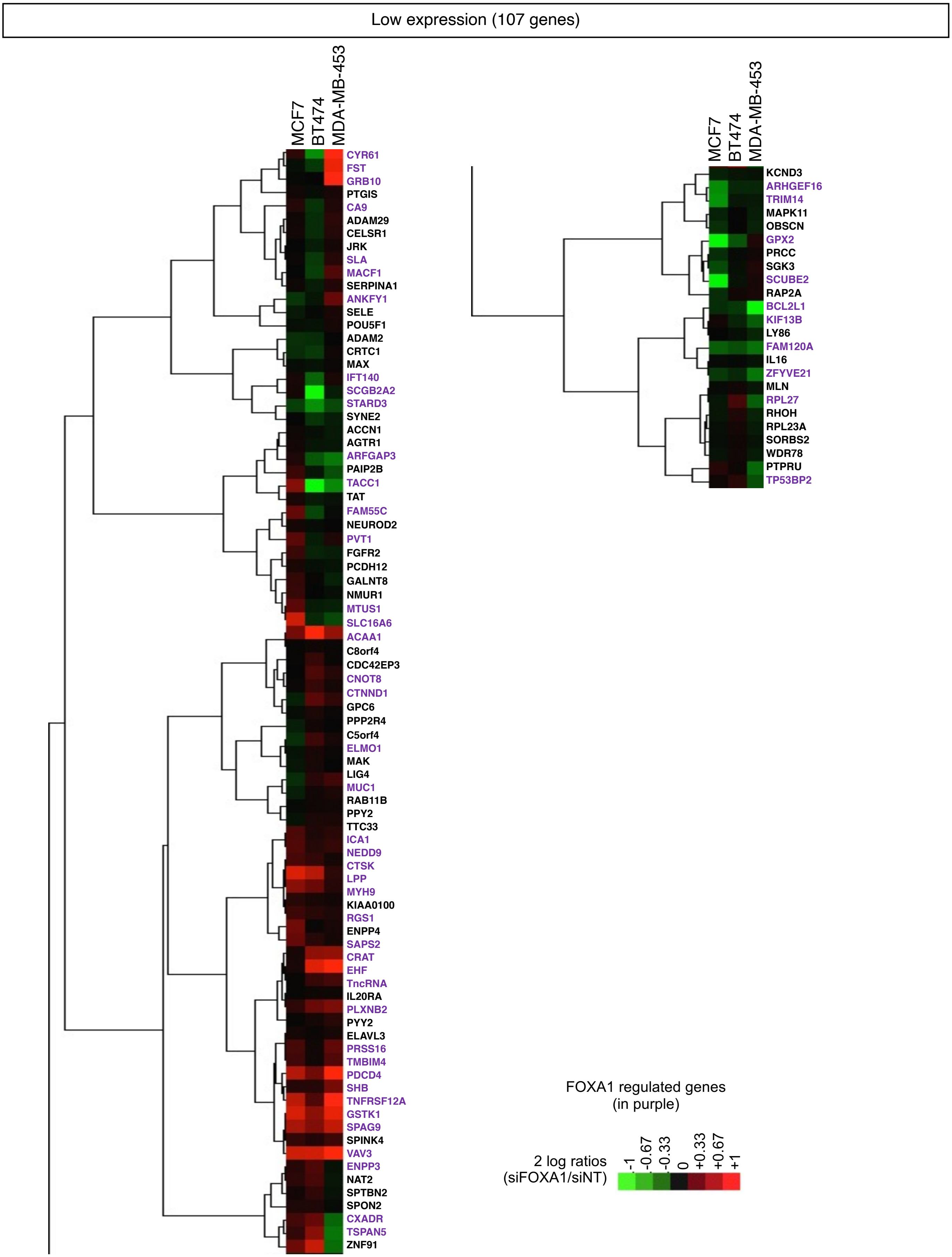

#### Figure Supplementary 6

GSEA of metastatic signature (vant Veer *et al*)

**A** The optimal set of 55 up-regulated genes predicting poor clinical outcome in breast cancer patients (defined as developing metastases with 5 years) from van't Veer *et al*)

| | |
|---|---|
| ALDH4A1 | MCM6 |
| AP2B1 | MELK |
| BBC3 | MMP9 |
| C16orf61 | MS4A7 |
| C17orf109 | MTDH |
| C20orf46 | NDC80 |
| CCNE2 | NMU |
| CDC42BPA | NUSAP1 |
| CENPA | ORC6 |
| COL4A2 | OXCT1 |
| DCK | PALM2-AKAP2 |
| DIAPH3 | PITRM1 |
| DTL | PRC1 |
| EBF4 | QSOX2 |
| ECI2 | RAB6B |
| ESM1 | RFC4 |
| EXT1 | RTN4RL1 |
| FGF18 | RUNDC1 |
| FLT1 | SCUBE2 |
| GMPS | SERF1A |
| GNAZ | SLC2A3 |
| GPR126 | STK32B |
| GPR180 | TGFB3 |
| GSTM3 | TSPYL5 |
| HRASLS | UCHL5 |
| IGFBP5 | WISP1 |
| LOC286052 | ZNF385B |
| LPCAT1 | |

**B** GSEA of FOXA1 up-regulated genes

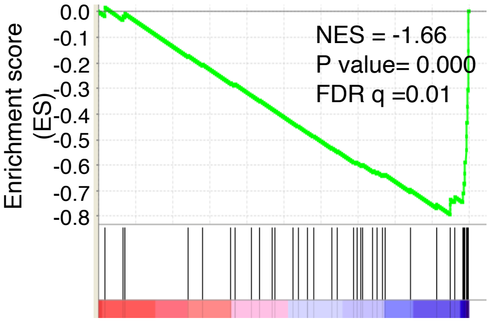


17/55 genes (37%)

| | |
|---|---|
| QSOX2 | CCNE2 |
| UCHL5 | MS4A7 |
| DCK | CENPA |
| DIAPH3 | NUSAP1 |
| MTDH | RFC4 |
| TGFB3 | MELK |
| RUNDC1 | NDC80 |
| TDL | MCM6 |
| | PRC1 |

**C** GSEA of FOXA1 down-regulated genes

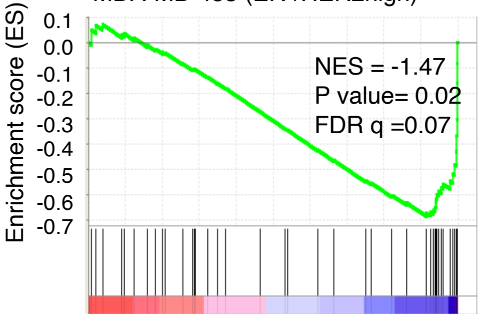


20/55 genes (43%)

| | |
|---|---|
| HRASLS | IGFBP5 |
| DCK | RUNDC1 |
| DIAPH3 | PRC1 |
| OXCT1 | MELK |
| UCHL5 | RFC4 |
| TGFB3 | CCNE2 |
| PITRM1 | DTL |
| CENPA | MCM6 |
| AP2B1 | STK32B |
| NUSAP1 | MS4A7 |

Shared genes between both HER2high cell lines → 14/55 genes (25%)

**D**

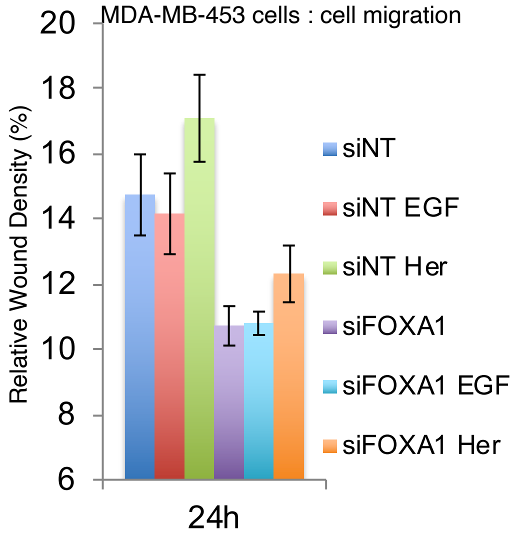

### Figure Supplementary 7

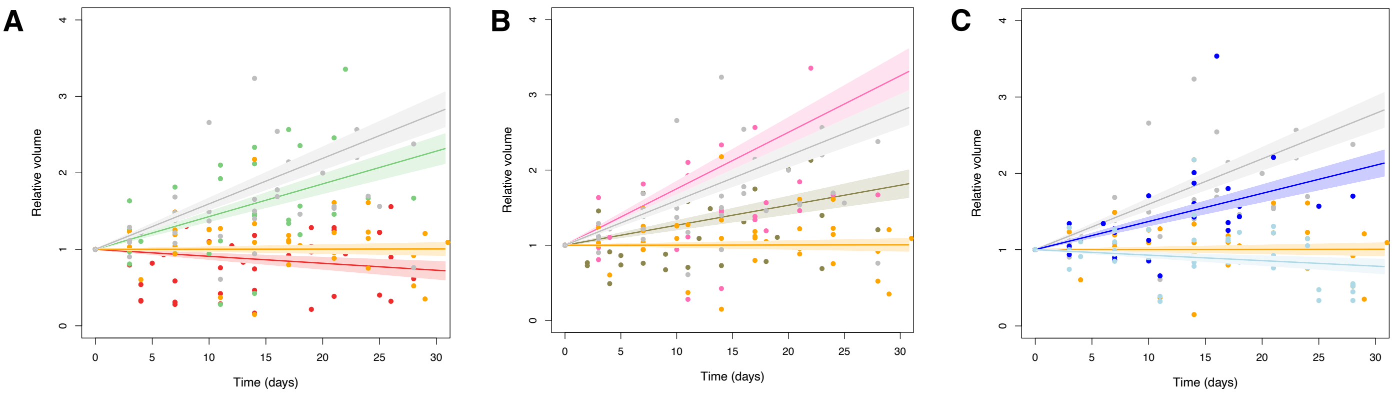

### Figure Supplementary 8

**A** ER levels in breast cancer cells

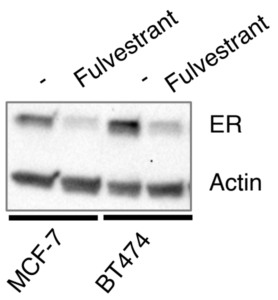


Cell growth: breast cancer cells

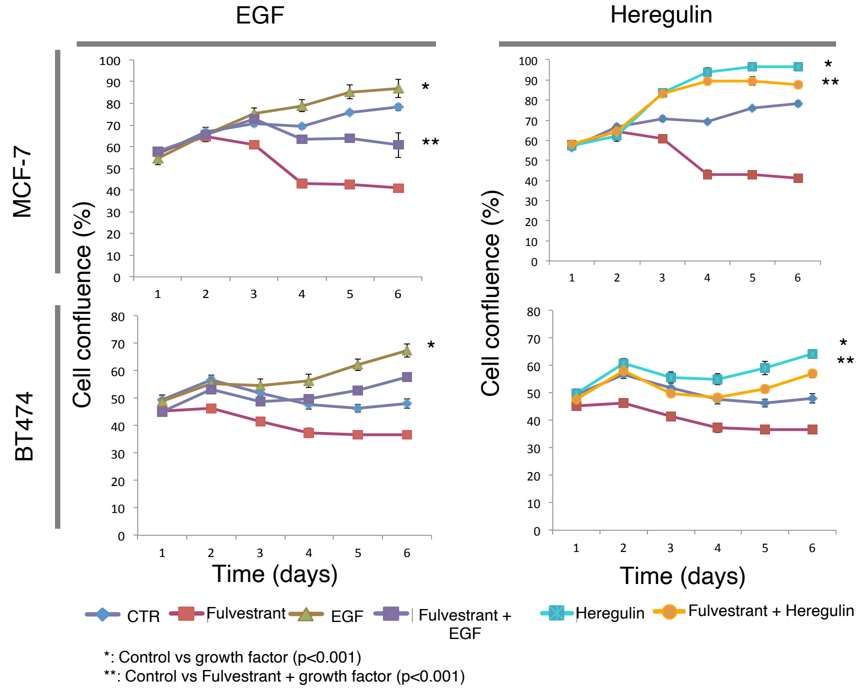


**B** ER and FOXA1 protein expression in PDX tumors

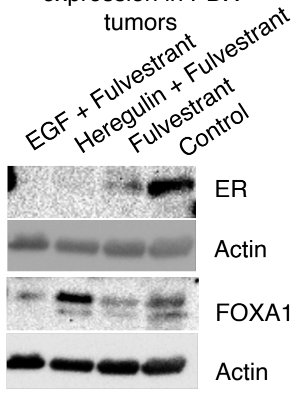


**C** FOXA1 ChIP-seq in PDX tumors (FOXA1 sites associated with poor prognosis)

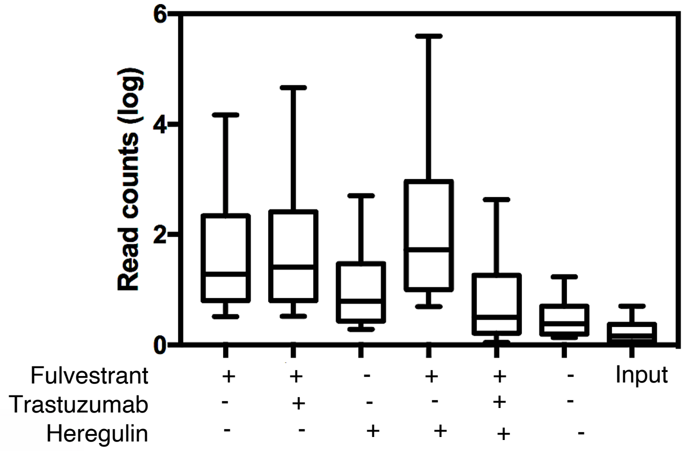


**D** Genomic distribution of FOXA1 sites at PDX tumors

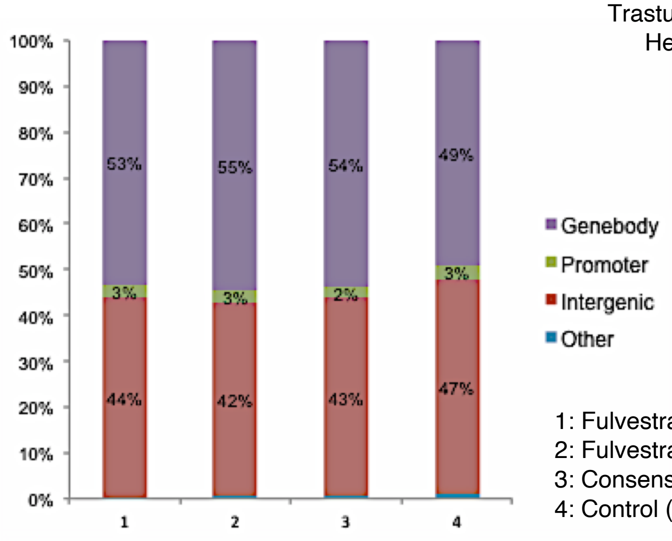


1: Fulvestrant and Hereglulin unique (4785 FOXA1 sites)
2: Fulvestrant unique (710 FOXA1 sites)
3: Consensus of 1 and 2 (62229 FOXA1 sites)
4: Control (257 FOXA1 sites)

### Figure Supplementary 9

**A** Genomic location of the three ERBB3 and the one FOXA1 mRNA probes analyzed for Metabric Project (Curtis *et al*)

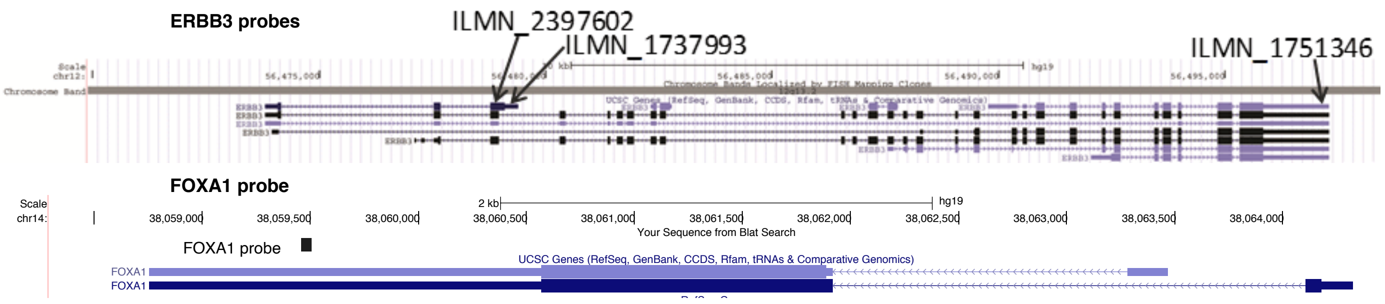


**B** Lineal regression of expression between FOXA1 and each of the 3 ERBB3 probes in Luminal A, Luminal B or HER2 breast cancer subtypes

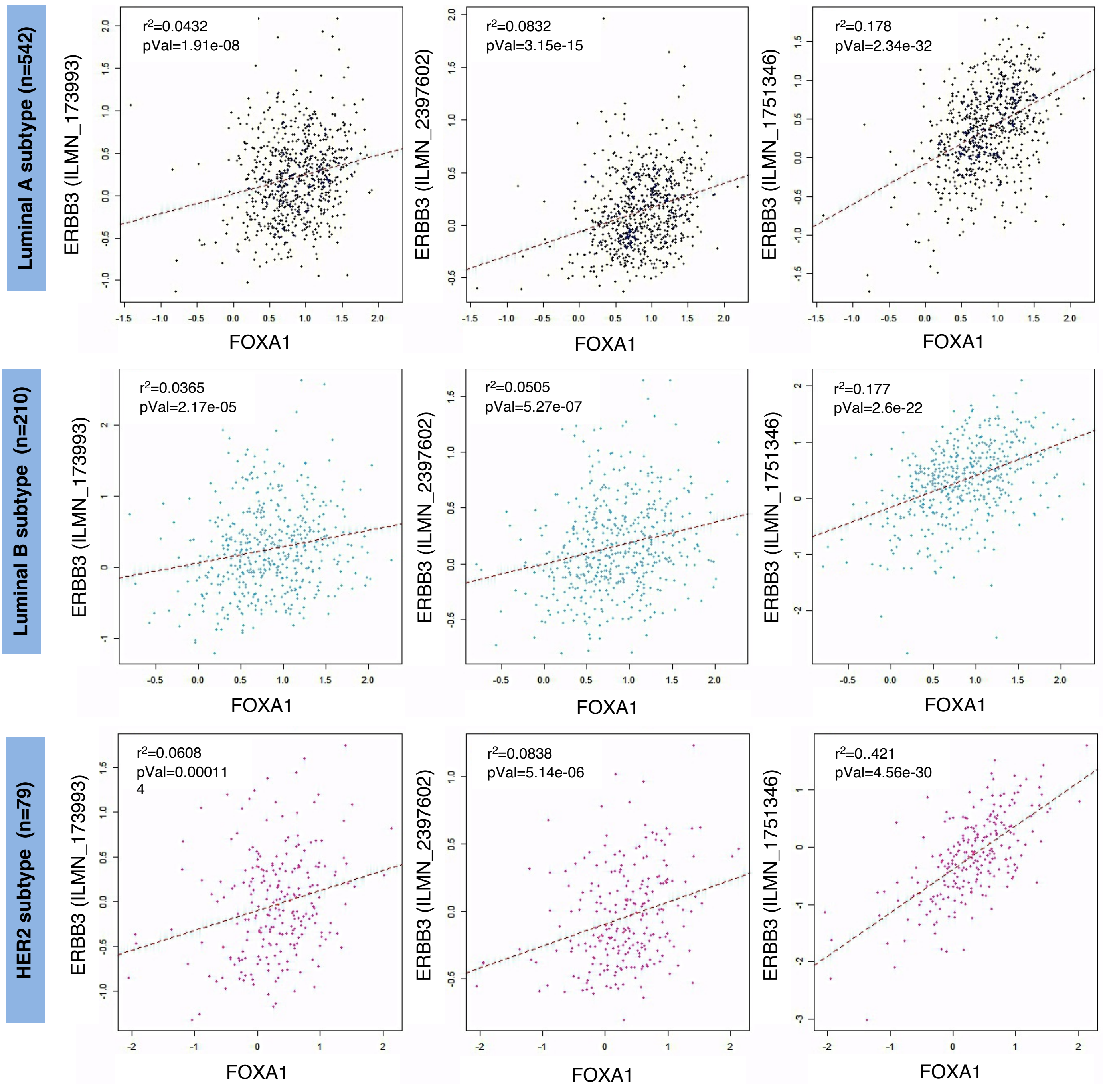

#### Figure Supplementary 10

**A** Motif analysis

| FOXA1 peaks | AP2Gamma 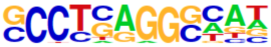 | PBX1 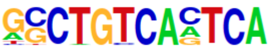 |
|---|---|---|
| MCF-specific | 6.32% (5.09%) p-value 1e-10 | 0.55% (0.50%) p-value 1 |
| MCF7-BT474 shared | 11.60% (5.39%) p-value 1e-513 | 0.58% (0.40%) p-value 1e-7 |
| BT474-specific | 14.41% (5.47%) p-value1e-1240 | 0.48% (0.37%) p-value 1e-4 |

**B** MCF-7 cells: FAIRE signal towards FOXA1 regulated genes (64% of the Heregulin induced regions and non-FOXA1 influenced)

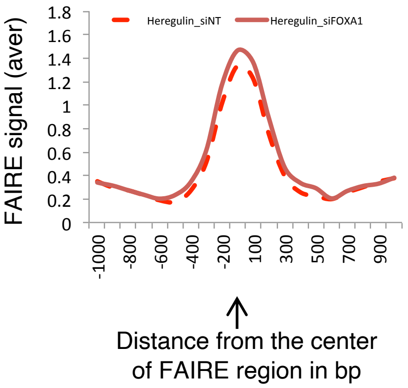


**C** FOXA1 binding at chromatin fraction

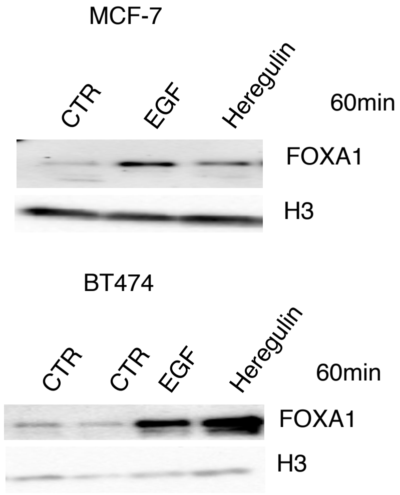


**D** FOXA1 ChIP upon Heregulin treatment

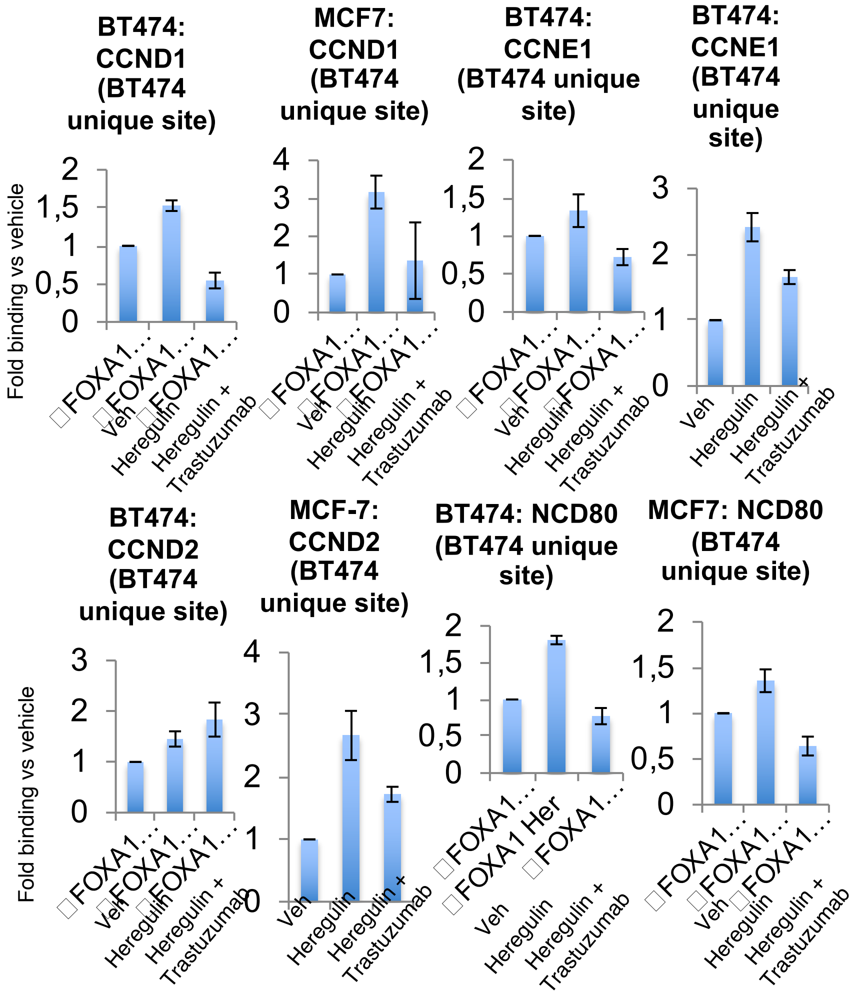

**Figure Supplementary 11**

**A** FOXA1 acetylation in MCF7 cells depleted with EP300

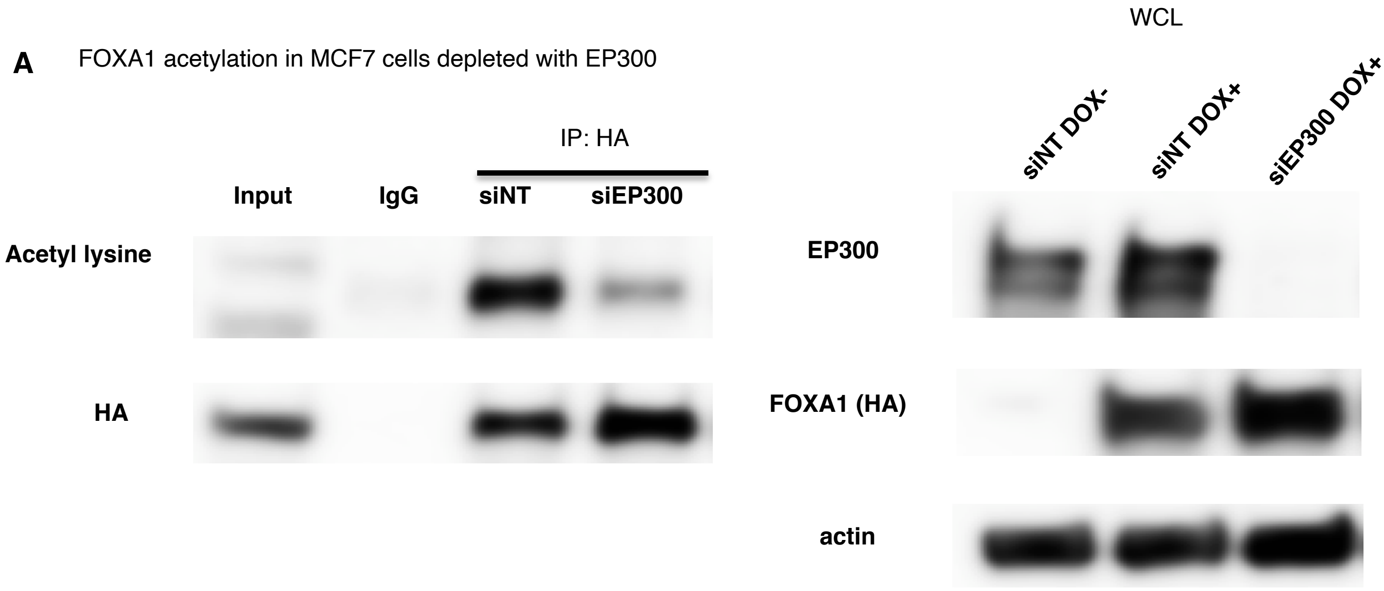


**B** Inhibition of HER2/HER3 signaling increased FOXA1 acetylation in MCF-7 HER2 high cell line

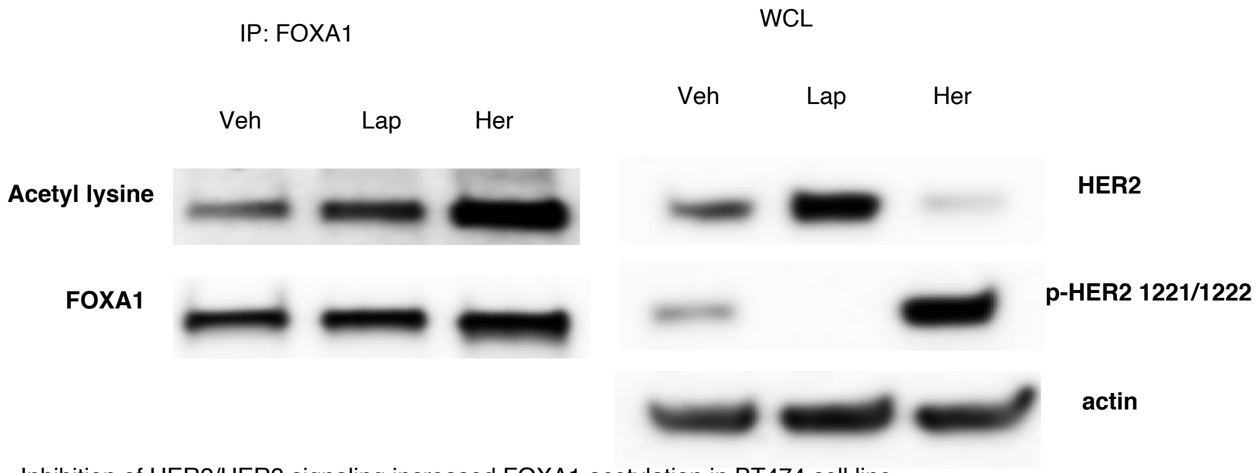


**C** Inhibition of HER2/HER3 signaling increased FOXA1 acetylation in BT474 cell line

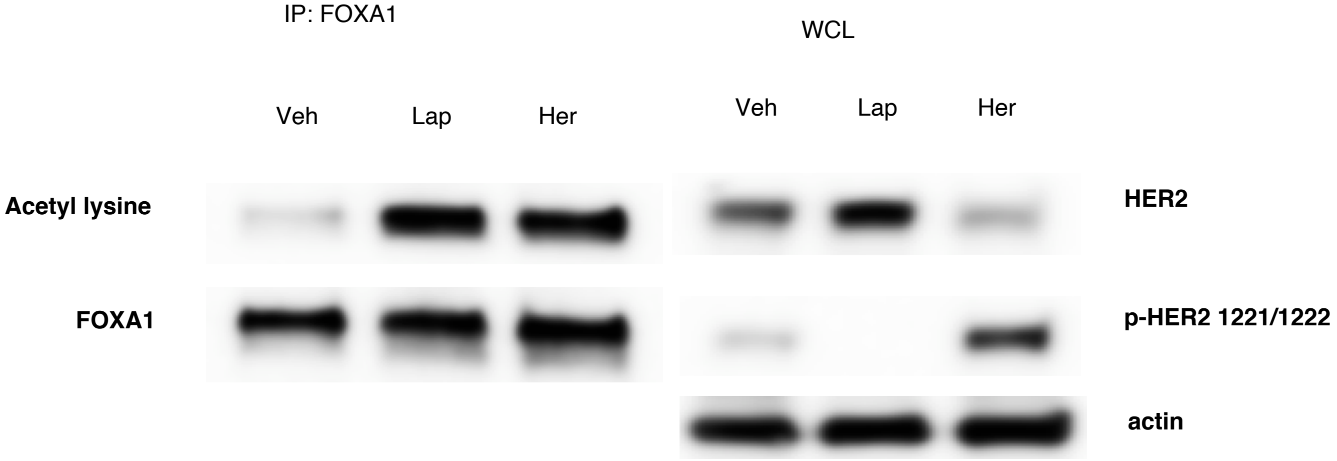

#### Supplementary Table 1

| | | N (%) of 15 patients |
|---|---|---|
| **Age (median, range)** | 43 (27-74) | |
| **Menopausal status** | Premenopausal | 10 (60) |
| | Postmenopausal | 5 (33) |
| | Unknown | 1 (7) |
| **Pathologic tumor size** | <2 cm | 13 (86) |
| | 2-5 cm | 1 (7) |
| | >5 cm | 0 |
| | Unknown | 1 (7) |
| **Number of involved axillary nodes** | 0 | 8 (53) |
| | 1-3 | 3 (20) |
| | ≥4 | 1 (7) |
| | Unknown | 3 (20) |
| **Histology** | IDC | 12 (80) |
| | ILC | 1 (6.66) |
| | Mixed (IDC+ILC) | 1 (6.66) |
| | Unknown | 1 (6.66) |
| **Histologic grade** | 1 | 12 (80) |
| | 2 | 1 (6.66) |
| | 3 | 1 (6.66) |
| | Unknown | 1 (6.66) |
| **Hormone Receptor Status** | ER+PR+ | 12 (75) |
| | ER+ PR- | 1 (12.5) |
| | ER- PR+ | 1 (6.2) |
| | Unknown | 1 (6.2) |
| **Adjuvant chemotherapy** | Yes | 12 (80) |
| | No | 2 (13) |

20 tumor samples from 15 patients were analyzed. 5 patients have paired tumor samples, from primary tumors and from relapse (metastases from patients after exposure to endocrine adjuvant therapies). Only characteristics at diagnoses of first primary tumors are described. In all patients the initial treatment for their primary tumors comprised either mastectomy (with/without radiation therapy) or lumpectomy plus radiation therapy, together with sentinel node biopsy and/or axillary dissection as clinically indicated.
ER: Estrogen Receptor, PR: Progesterone Receptor, IDC: Invasive Ductal carcinoma, ILC: Invasive Lobular Carcinoma, AI: Aromatase Inhibitor.
ER and PR positivity was defined as >1% nuclei stained. HER2 positivity was defined according to 2007 American College of Pathologist Guidelines.

#### Supplementary Table 2

Linear regression parameters for each treatment group (1: active substance, 0: vehicle).

| Fulvestrant | Heregulin | Trastuzumab | EGF | beta.pvalue | beta | beta.se | rvolume20.pred | rvolume20.se |
|---|---|---|---|---|---|---|---|---|
| 1 | 1 | 1 | 0 | 0,020 | -0,009 | 0,004 | 0,818 | 0,075 |
| 0 | 1 | 1 | 0 | 2,5E-09 | 0,029 | 0,004 | 1,587 | 0,076 |
| 1 | 0 | 1 | 0 | 0,311 | 0,003 | 0,003 | 1,062 | 0,061 |
| 0 | 0 | 1 | 0 | 6,1E-05 | 0,027 | 0,006 | 1,532 | 0,117 |
| 1 | 1 | 0 | 0 | 4,2E-08 | 0,043 | 0,006 | 1,856 | 0,125 |
| 0 | 1 | 0 | 0 | 7,5E-09 | 0,075 | 0,010 | 2,503 | 0,194 |
| 1 | 0 | 0 | 0 | 0,953 | 1,6E-04 | 0,003 | 1,003 | 0,053 |
| 0 | 0 | 0 | 0 | 2,5E-10 | 0,059 | 0,007 | 2,190 | 0,146 |
| 1 | 0 | 0 | 1 | 0,020 | -0,007 | 0,003 | 0,856 | 0,060 |
| 0 | 0 | 0 | 1 | 8,0E-08 | 0,037 | 0,005 | 1,737 | 0,109 |
